# Varied forms of mutant tRNA mistranslation produce distinct phenotypes across multiple model organisms

**DOI:** 10.64898/2026.08.25.747118

**Authors:** Brendan S. Charles, Amanda J. Moehring

## Abstract

Mutant tRNA mistranslation is a phenomenon in which specific tRNA gene mutations cause translational errors wherein amino acids incorporated during translation differ from those coded by mRNA sequences. The biological consequences of mutant tRNA mistranslation are highly complex and contextual, but are often deleterious. Importantly, many variables exist that are likely to affect the outcomes of any one mutant mistranslating tRNA; biochemical characteristics of the exchanged amino acids, frequency of use, identity of mistranslated products, and so on. Here, we generate a carefully curated array of tRNA mutants to assess which characteristics of mistranslating tRNA variants are most predictive of deleterious phenotypes. We find that toxicity often arises from mutant tRNAs that induce dramatic changes in biochemical properties between exchanged amino acids, as well as mistranslation that occurs more frequently. Importantly, exceptions are also observed to each of these general rules, implying instead that some deliriousness may arise from more granular, product-specific mechanisms. Additionally, quantification of mistranslation via mass spectrometry demonstrates a weak relationship between quantity of mistranslated products and severity of toxic phenotypes. Lastly, despite previous establishment of mutant mistranslating tRNA models in *Drosophila melanogaster*, incorporation of higher frequency mistranslating tRNA variants was largely unsuccessful, and incorporation of lower frequency mistranslating tRNA variants produces no detectable developmental phenotypes. These experiments support the notion that although general rules may be capable of reasonably predicting their consequences, each mistranslating tRNA variant warrants individual consideration and investigation for thorough understanding of its biological outcomes and precise mechanisms of toxicity.

## Introduction

Translational error, or mistranslation, results in nascent proteins incorporating amino acids that differ from those coded by mRNA transcripts (Moghal et al., 2014; Zimmerman et al., 2018). The protein products of mistranslation are therefore mutants, despite originating from non-mutant mRNA sequences. These mistranslated mutant proteins, like other mutant proteins, may be prone to misfolding, thus increasing load on proteostatic mechanisms. Mistranslation occurs naturally at low frequencies and is generally well tolerated (Berg & Brandl, 2021; Joshi et al., 2019; Loftfield & Vanderjagt, 1972; Moghal et al., 2014), likely due to the multitude of mechanisms by which cells can attenuate proteostatic stresses (Labbadia & Morimoto, 2015; McDonald et al., 2025; Roth & Balch, 2011). However, various mutations to translational machinery, such as tRNA genes or the aminoacyl tRNA synthetases (aaRSs) that charge tRNAs with their corresponding amino acid, can greatly elevate the frequency of mistranslation (Cozma et al., 2023; Davey-Young et al., 2024; Lee et al., 2006; Liu et al., 2014; Zimmerman et al., 2018).

tRNAs bearing specific mutations can greatly increase the frequency of mistranslation such that deleterious phenotypes of varying severity arise. Eukaryotic cell models expressing mistranslating tRNA mutants regularly display growth deficits, demonstrating a toxic effect for elevated mistranslation (Berg et al., 2017; Cozma et al., 2023; Davey-Young et al., 2024; Hasan et al., 2023; Lant et al., 2018, 2021; Zimmerman et al., 2018). The induction of proteostatic stress response pathways, namely heat shock response and autophagy, have also been observed in response to tRNA mistranslation (Berg et al., 2021; Hasan et al., 2023; McDonald et al., 2025). Activation of stress response pathways of these varieties implicate proteotoxicity as a mechanism for eliciting arrested growth in models of mutant tRNA mistranslation. Our previous work in multicellular models demonstrates that tRNA mistranslation can cause both increased developmental lethality as well as more frequent display of developmental deformities (Isaacson et al., 2022, 2024).

Not all mutant tRNA mistranslation is equal in its biological outcomes. Some mistranslating tRNAs display no such negative phenotypes and appear benign (Cozma et al., 2023; Davey-Young et al., 2024; Hasan et al., 2023; Zimmerman et al., 2018). Mistranslation of specific varieties has even been shown to be contextually adaptive (Fan et al., 2015; Lee et al., 2014; Schwartz & Pan, 2016). In spite of the aforementioned developmental lethality in multicellular models, individuals that survive into adulthood exhibit increased lifespan (Branco et al., 2026; Isaacson et al., 2022, 2024), demonstrating a circumstantial aspect of the deleterious effects of some mistranslation.

A plethora of variables are likely to contribute to the diverse biological outcomes caused by mutant tRNA mistranslation. First, the biochemical differences between the substituted amino acids is likely relevant to the propensity for mistranslated proteins to misfold, and thus the biological outcomes of their increase in prevalence. Additionally, mutant mistranslating tRNAs may differ in how frequently they induce mistranslation (Berg et al., 2017; Hummel et al., 2019). Given that different amino acids are used at different frequencies throughout the proteome (Sabbía et al., 2007), mistranslation of more common amino acids stand to create more proteostatic load, whereas mistranslation of rare amino acids may be comparatively well tolerated. There are also frequency of use differences at the codon level; despite several codons often encoding the same amino acid, not all codons within such a group are used equally in the transcriptome. This codon bias, referred to as codon optimality (Cochella & Green, 2005; Endres et al., 2015; Grantham et al., 1980; Komar, 2021; Quax et al., 2015; Saunders & Deane, 2010), has important implications for translational dynamics and efficiency of specific transcripts. As a result, mistranslation of particular codons may confer more deleterious effects than mistranslation of synonymous codons.

Further complicating the notion of varied mistranslation frequency is the concept of tRNA pool competition (Dayanand et al., 2018; Gao et al., 2024; Goodarzi et al., 2016; Sagi et al., 2016; Torres et al., 2019). Not all anticodons are equally represented within the tRNA pool. Therefore, the effects of mutant tRNA mistranslation may also be modulated by the volume of similar tRNAs competing for the same codons.

To investigate which characteristics of mutant tRNA mistranslation inform biological outcomes, we created an array of mutant tRNAs which mistranslate various codons and transformed them into both yeast and flies (Table 1). The majority of mistranslating mutants assessed here are variants of tRNA^Ser^, which was chosen for its ability to retain serine aminoacylation despite any anticodon mutation, therefore allowing the assessment of many forms of mistranslation from one tRNA gene. Additionally, tRNA^Ser^ has been used effectively in both of the model organisms used here, therefore minimizing potential troubleshooting risks. The generated array of mutants was carefully designed to address key characteristics of mistranslation, such as the severity of biochemical difference in substituted amino acids, anticipated frequency of codon use, and competition with similar tRNAs. Each mutant mistranslating tRNA represents a unique combination of each of these characteristics, or “form” of mistranslation. The effects of each form of mistranslation on general measures of model organism health are then used to inform which characteristics of mistranslation are especially relevant to biological outcomes. We hypothesized that mistranslation producing more substantial biochemical changes in the exchanged amino acids would result in more severe growth phenotypes, as would forms of mistranslation that occur more frequently.

**Table 1.**
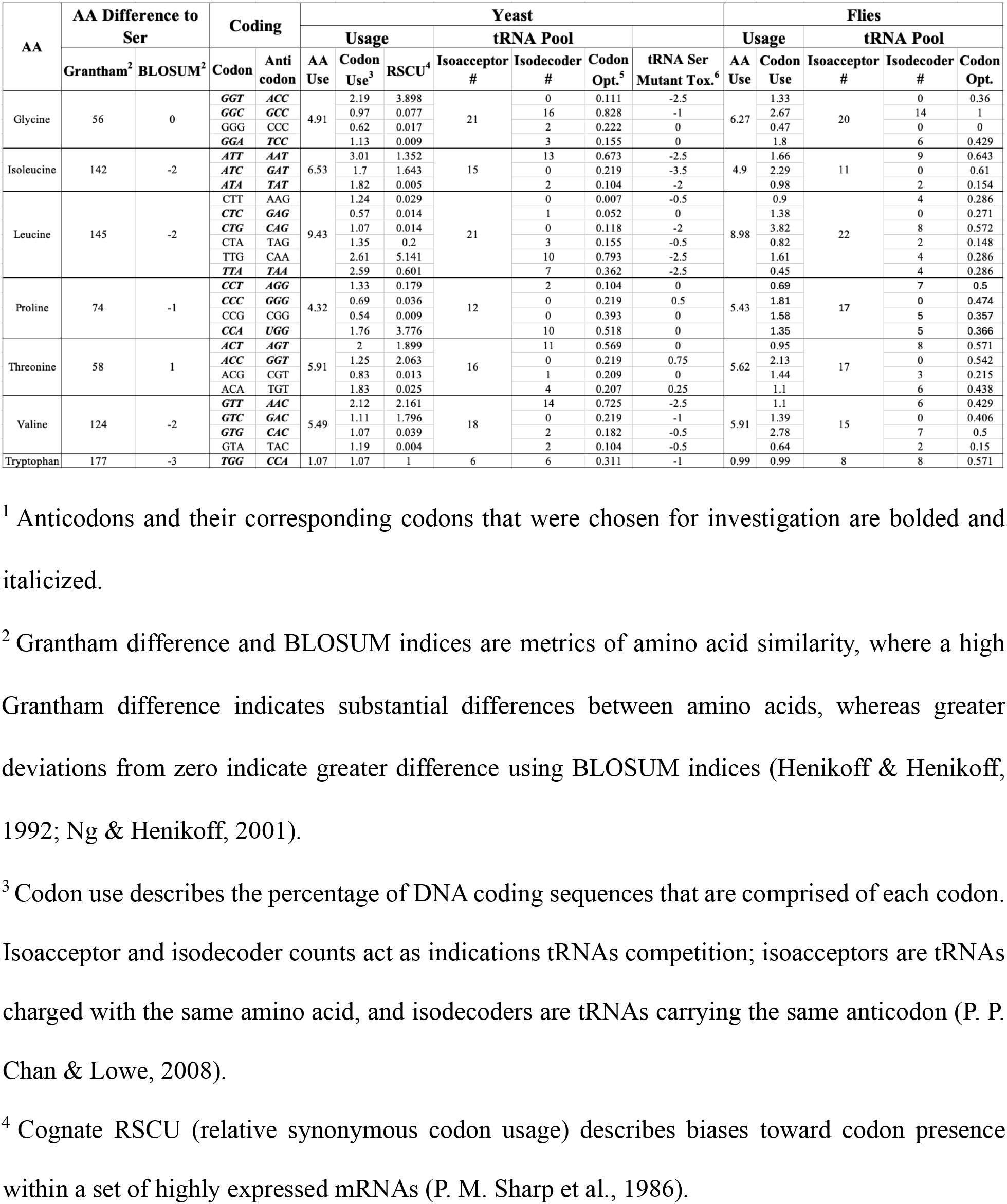
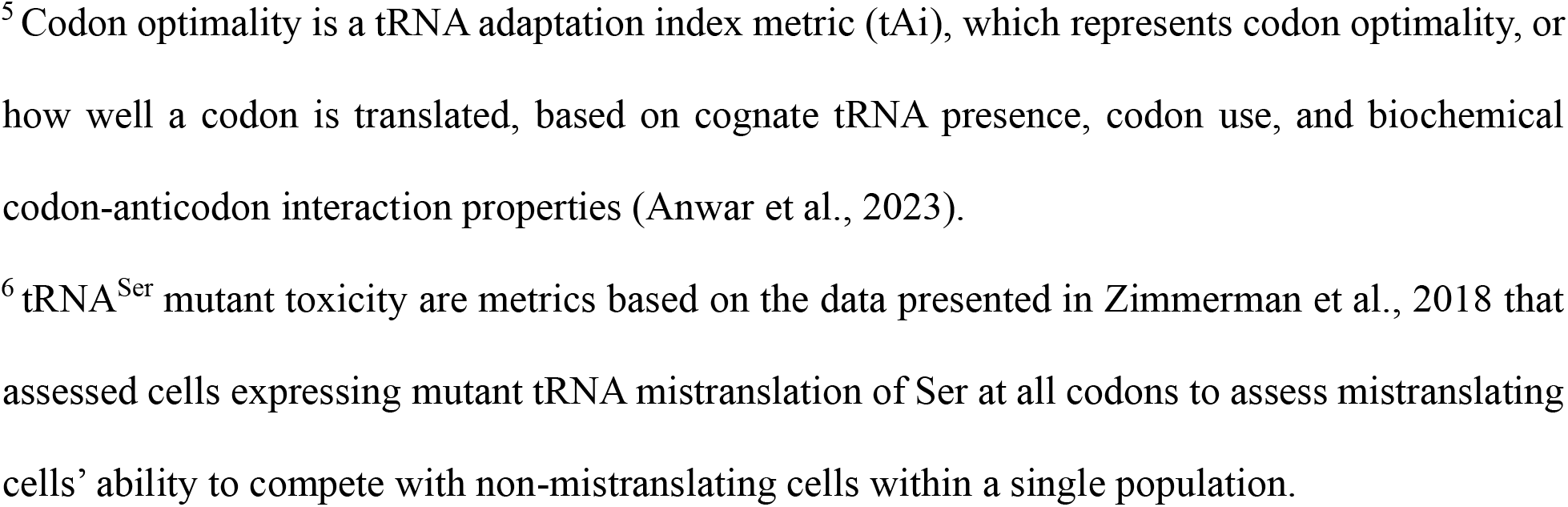
Variables considered for the choice of tRNA^Ser^ mistranslating anticodon mutants to be expressed in both yeast and *Drosophila melanogaster*.

| AA | AA Difference to Ser |  | Coding |  | Yeast |  |  |  |  |  |  | Flies |  |  |  |  |
|---|---|---|---|---|---|---|---|---|---|---|---|---|---|---|---|---|
|  |  |  |  |  | Usage |  |  | tRNA Pool |  |  |  | Usage |  | tRNA Pool |  |  |
|  | Grantham<sup>2</sup> | BLOSUM<sup>2</sup> | Codon | Anti codon | AA Use | Codon Use<sup>3</sup> | RSCU<sup>4</sup> | Isoacceptor # | Isodecoder # | Codon Opt.<sup>5</sup> | tRNA Ser Mutant Tox.<sup>6</sup> | AA Use | Codon Use | Isoacceptor # | Isodecoder # | Codon Opt. |
| Glycine | 56 | 0 | ***GGT*** | ***ACC*** | 4.91 | 2.19 | 3.898 | 21 | 0 | 0.111 | -2.5 | 6.27 | 1.33 | 20 | 0 | 0.36 |
|  |  |  | ***GGC*** | ***GCC*** |  | 0.97 | 0.077 |  | 16 | 0.828 | -1 |  | 2.67 |  | 14 | 1 |
|  |  |  | GGG | CCC |  | 0.62 | 0.017 |  | 2 | 0.222 | 0 |  | 0.47 |  | 0 | 0 |
|  |  |  | ***GGA*** | ***TCC*** |  | 1.13 | 0.009 |  | 3 | 0.155 | 0 |  | 1.8 |  | 6 | 0.429 |
| Isoleucine | 142 | -2 | ***ATT*** | ***AAT*** | 6.53 | 3.01 | 1.352 | 15 | 13 | 0.673 | -2.5 | 4.9 | 1.66 | 11 | 9 | 0.643 |
|  |  |  | ***ATC*** | ***GAT*** |  | 1.7 | 1.643 |  | 0 | 0.219 | -3.5 |  | 2.29 |  | 0 | 0.61 |
|  |  |  | ***ATA*** | ***TAT*** |  | 1.82 | 0.005 |  | 2 | 0.104 | -2 |  | 0.98 |  | 2 | 0.154 |
| Leucine | 145 | -2 | CTT | AAG | 9.43 | 1.24 | 0.029 | 21 | 0 | 0.007 | -0.5 | 8.98 | 0.9 | 22 | 4 | 0.286 |
|  |  |  | ***CTC*** | ***GAG*** |  | 0.57 | 0.014 |  | 1 | 0.052 | 0 |  | 1.38 |  | 0 | 0.271 |
|  |  |  | ***CTG*** | ***CAG*** |  | 1.07 | 0.014 |  | 0 | 0.118 | -2 |  | 3.82 |  | 8 | 0.572 |
|  |  |  | CTA | TAG |  | 1.35 | 0.2 |  | 3 | 0.155 | -0.5 |  | 0.82 |  | 2 | 0.148 |
|  |  |  | TTG | CAA |  | 2.61 | 5.141 |  | 10 | 0.793 | -2.5 |  | 1.61 |  | 4 | 0.286 |
|  |  |  | ***TTA*** | ***TAA*** |  | 2.59 | 0.601 |  | 7 | 0.362 | -2.5 |  | 0.45 |  | 4 | 0.286 |
| Proline | 74 | -1 | ***CCT*** | ***AGG*** | 4.32 | 1.33 | 0.179 | 12 | 2 | 0.104 | 0 | 5.43 | 0.69 | 17 | 7 | 0.5 |
|  |  |  | ***CCC*** | ***GGG*** |  | 0.69 | 0.036 |  | 0 | 0.219 | 0.5 |  | 1.81 |  | 0 | 0.474 |
|  |  |  | CCG | CGG |  | 0.54 | 0.009 |  | 0 | 0.393 | 0 |  | 1.58 |  | 5 | 0.357 |
|  |  |  | ***CCA*** | ***UGG*** |  | 1.76 | 3.776 |  | 10 | 0.518 | 0 |  | 1.35 |  | 5 | 0.366 |
| Threonine | 58 | 1 | ***ACT*** | ***AGT*** | 5.91 | 2 | 1.899 | 16 | 11 | 0.569 | 0 | 5.62 | 0.95 | 17 | 8 | 0.571 |
|  |  |  | ***ACC*** | ***GGT*** |  | 1.25 | 2.063 |  | 0 | 0.219 | 0.75 |  | 2.13 |  | 0 | 0.542 |
|  |  |  | ACG | CGT |  | 0.83 | 0.013 |  | 1 | 0.209 | 0 |  | 1.44 |  | 3 | 0.215 |
|  |  |  | ACA | TGT |  | 1.83 | 0.025 |  | 4 | 0.207 | 0.25 |  | 1.1 |  | 6 | 0.438 |
| Valine | 124 | -2 | ***GTT*** | ***AAC*** | 5.49 | 2.12 | 2.161 | 18 | 14 | 0.725 | -2.5 | 5.91 | 1.1 | 15 | 6 | 0.429 |
|  |  |  | ***GTC*** | ***GAC*** |  | 1.11 | 1.796 |  | 0 | 0.219 | -1 |  | 1.39 |  | 0 | 0.406 |
|  |  |  | ***GTG*** | ***CAC*** |  | 1.07 | 0.039 |  | 2 | 0.182 | -0.5 |  | 2.78 |  | 7 | 0.5 |
|  |  |  | GTA | TAC |  | 1.19 | 0.004 |  | 2 | 0.104 | -0.5 |  | 0.64 |  | 2 | 0.15 |
| Tryptophan | 177 | -3 | ***TGG*** | ***CCA*** | 1.07 | 1.07 | 1 | 6 | 6 | 0.311 | -1 | 0.99 | 0.99 | 8 | 8 | 0.571 |

<sup>1</sup> Anticodons and their corresponding codons that were chosen for investigation are bolded and italicized.
<sup>2</sup> Grantham difference and BLOSUM indices are metrics of amino acid similarity, where a high Grantham difference indicates substantial differences between amino acids, whereas greater deviations from zero indicate greater difference using BLOSUM indices (Henikoff & Henikoff, 1992; Ng & Henikoff, 2001).
<sup>3</sup> Codon use describes the percentage of DNA coding sequences that are comprised of each codon. Isoacceptor and isodecoder counts act as indications tRNAs competition; isoacceptors are tRNAs charged with the same amino acid, and isodecoders are tRNAs carrying the same anticodon (P. P. Chan & Lowe, 2008).
<sup>4</sup> Cognate RSCU (relative synonymous codon usage) describes biases toward codon presence within a set of highly expressed mRNAs (P. M. Sharp et al., 1986).
<sup>5</sup> Codon optimality is a tRNA adaptation index metric (tAi), which represents codon optimality, or how well a codon is translated, based on cognate tRNA presence, codon use, and biochemical codon-anticodon interaction properties (Anwar et al., 2023).
<sup>6</sup> tRNA<sup>Ser</sup> mutant toxicity are metrics based on the data presented in Zimmerman et al., 2018 that assessed cells expressing mutant tRNA mistranslation of Ser at all codons to assess mistranslating cells' ability to compete with non-mistranslating cells within a single population.

To address each of our research questions, we used the model organisms *Saccharomyces cerevisiae* and *Drosophila melanogaster*, more commonly known as baker’s yeast and fruit flies, respectively. These models were chosen for their tractability in the integration and assessment of several different genetic variants – compared to other models, the creation of novel transgenic models can be accomplished faster and with greater ease. Similarly, assessments of general health metrics and other phenotypes can be carried out relatively quickly. While yeast cells are one of the most common models in which to study mutant tRNA mistranslation and are suitable for probing many variants in tandem, comparatively little research has been done assessing mutant tRNA mistranslation in multicellular organisms. Here, the use of fruit flies, a multicellular organism with abundant genetic tools and deeply characterized physiological and behavioural phenotypes, aims to bridge this research gap.

First, to test how severity of biochemical difference in exchanged amino acids and codon use influence cell health, our array of mutant mistranslating tRNAs are tested for growth rate in yeast cells. Each of these varied forms of mistranslation are also assessed for their ability to suppress growth deficits imbued by mistranslation when permitted to mistranslate for longer time periods. Next, the influence of mistranslation frequency on phenotype is assessed in two ways: variable rate of mistranslation and the use of alternative uniquely expressed tRNA genes bearing the same mistranslating mutation. The rate of mistranslation was manipulated via the addition of secondary mutations which reduce mutant tRNA participation in translation. The role of specific codon identity in mediating outcomes of mistranslation was assessed by measuring yeast cell growth changes stemming from the mistranslation of high frequency optimal codons compared to more rare, low-optimality codon mistranslation for the same amino acid substitution. Mistranslating mutations to tRNA genes of different amino acid isoacceptor families were also assessed to probe alternative forms of mistranslation.

To ensure the observed phenotypes in cells expressing these mutant tRNAs can confidently be attributed to mistranslation, confirmation and quantification of mistranslation was accomplished via mass spectrometry. Lastly, to assess whether the biological impacts of these various forms of mistranslation are conserved in a multicellular model, their integration into *Drosophila melanogaster* was attempted. Of the few viable lines created, health effects of mistranslation were assessed in *Drosophila* by developmental viability screening, then compared to the effects of the same mistranslating tRNAs in yeast. The data generated by these experiments demonstrate that the severity of biochemical difference in exchanged amino acids is a key variable in predicting the deleterious effects of mistranslation, with mistranslation frequency also acting as a key factor.

## Methods

### tRNA mutant design

In an effort to determine patterns of which forms of mistranslating tRNA variants are deleterious, and how their phenotypes differ between model organisms, we designed an array of anticodon mutants with which to assess different aspects of mistranslation, namely severity of biochemical differences between substituted amino acids and frequency of cognate codon use. These anticodons and their various characteristics are described in Table 1, and were all generated from the *D. melanogaster* tSerUGA1-1 gene (Ser denotes isoacceptor group, UGA denotes isodecoder group, 1-1 denotes isoform group and specific genomic copy). Amino acid differences were determined by BLOSUM matrices and as well as Grantham Differences (Grantham, 1974; Henikoff & Henikoff, 1992; Ng & Henikoff, 2001).

Pro-UGG was chosen for its use in previous research as a means of comparison (Berg et al., 2017). To assess the influence of low biochemical difference in exchanged amino acids, we selected Thr-AGT as threonine represents the most similar AA to serine. The AGT codon chosen for Thr was selected for its predicted frequency of use being similar to that of other chosen codons, based on cognate codon prevalence, competitor tRNA counts, and codon optimality. Glycine represents a relatively conservative biochemical difference but is particularly abundant in NDD associated proteins (Table S1). The frequency of use for the chosen Gly-ACC anticodon is also predicted to be similar to that of the codons above, due to a high prevalence of cognate codons but substantial competitor tRNA pool. Valine is similarly abundant in NDD associated proteins (Table S1) but represents a more substantial biochemical difference from serine than either glycine or proline. Thus, the use of Val-ACC and Gly-AAC allow probing of whether more frequent mistranslation of codons more represented among NDD proteins results in greater toxicity when both mistranslating mutant tRNAs and NDD proteins are expressed together. The Val-ACC anticodon was also chosen for similarities in the aforementioned frequency of use metrics. Trp-CCA, Ile-AAT, and Leu-TAA were all chosen for their substantial biochemical differences from serine, while also differing between one another in their frequency of use; tryptophan is rare, while Ile-AAT and Leu-TAA are common. Variables considered for mutant design are outlined in Table 1. An expanded version of Table 1 exploring all anticodons can be found in Table S2.

An additional important factor contributing to mistranslation frequency is the transcriptional activity of the mutant tRNA. As such, we set out to assess whether mistranslation toxicity changes with different tRNA^Ser^ genes bearing the same mistranslating mutations. Table 2 contains the information used to choose *D. melanogaster* tRNA^Ser^ genes to mutagenize. tSerAGA1-1 was chosen for its low chromatin accessibility score as determined by ATAC Seq (Merrill et al., 2022; Milon et al., 2014), which we hypothesize corresponds to low expression. tSerAGA2-4 is situated among highly open chromatin, which we hypothesize to correspond to higher expression, and has the same competitor isodecoder pool as tSerAGA1-1. tSerGCT2-2 and tSerGCT2-4 were also attempted but could not be PCR amplified from genomic DNA. Each of the chosen tRNA^Ser^ isoforms was mutated to contain Pro-UGG and Gly-ACC anticodons, shown in Figure 1 to be well tolerated or toxic, respectively, when expressed in tSerUGA1-1.

**Figure 1.**
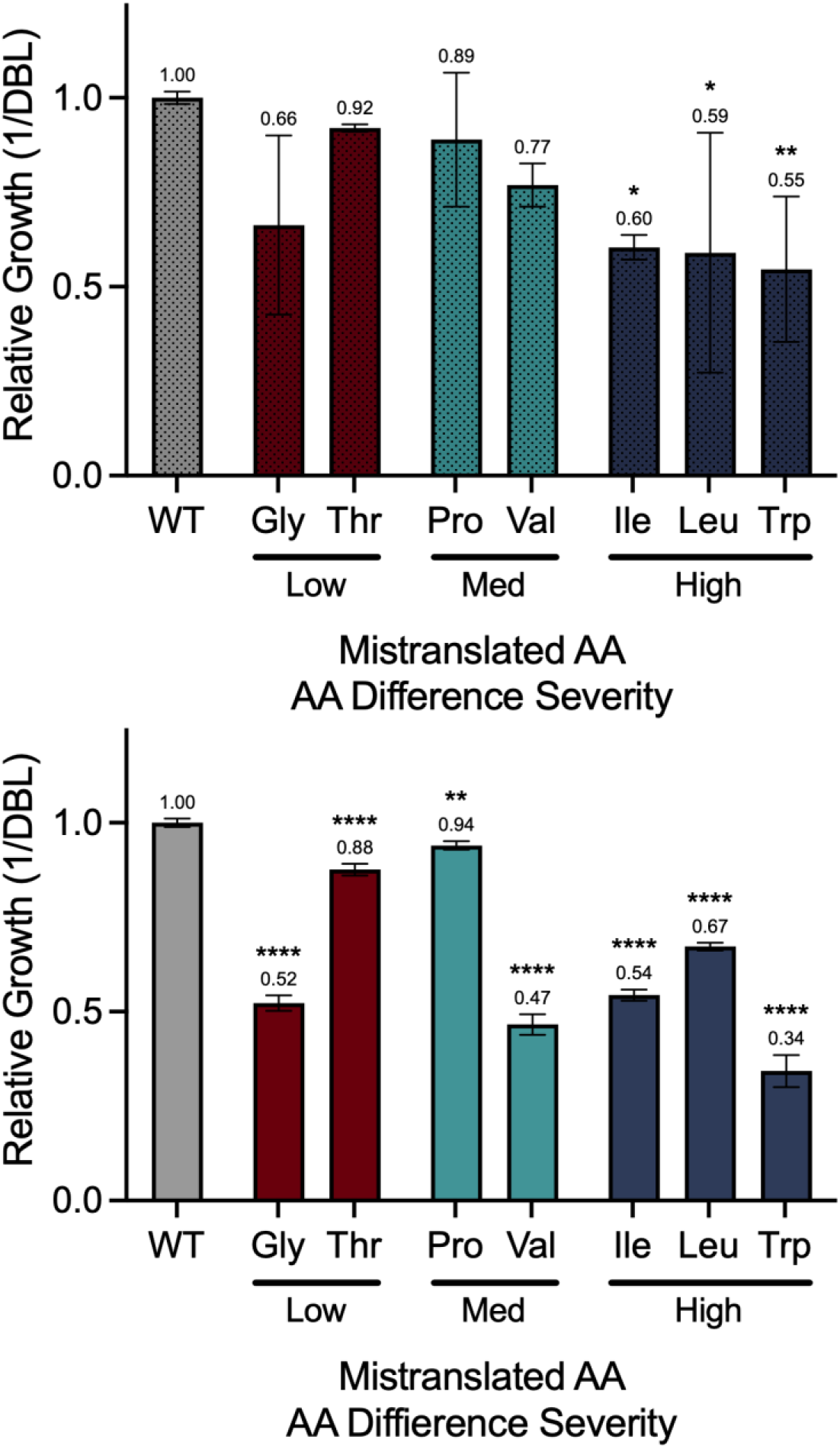
Mistranslation induced growth deficits are affected by both form of mistranslation and time spent mistranslating. Forms of mistranslation are grouped by colour according to qualitative magnitude of biochemical difference to serine: Gly and Thr are a low magnitude of change (maroon), Pro and Val comprise a medium magnitude of change (teal), whereas Ile, Leu, and Trp are a high magnitude of change (navy). Top: constitutive expression of mistranslating tRNA variants mistranslating Ser at Gly-ACC, Ile-ATT, Leu-TAA, Pro-UGG, Thr-AGT, Trp-CCA, and Val-AAC. Bottom: expression of the same tRNA variants limited exclusively to the 24-hour growth assay via inducible tRNA expression system. Group means are listed as annotations above each bar (n=4). Error bars denote standard deviation. Statistical significance as determined by one-way ANOVA and subsequent post-hoc pairwise Dunnet’s multiple comparisons tests is marked by asterisks (**** denotes p<0.00001, ** denotes p<0.01, * denotes p<0.05). Growth curves can be found in Supplementary Figure S1.

**Table 2.** Characteristics of alternative tRNA genes used to plan mutagenesis.

| **tRNA gene<sup>1</sup>** | **ATACSeq<sup>2</sup>** | **Isoacceptor #<sup>3</sup>** | **Isodecoder #<sup>4</sup>** | **Cognate RSCU<sup>5</sup>** | **Codon Optimality<sup>6</sup>** |
|---|---|---|---|---|---|
| tSerUGA1-1 | 845 | 19 | 3 | 0.122 | 0.155 |
| tSerAGA2-4 | 655 | 19 | 11 | 3.359 | 0.236 |
| tSerAGA1-1* | 0 | 19 | 11 | 3.359 | 0.236 |
| tSerGCU2-2* | 23 | 19 | 4 | 0.105 | 0.104 |
| tSerGCU2-4* | 535 | 19 | 4 | 0.105 | 0.104 |
| tAlaAGC2-1* | 143 | 16 | 11 | 3.005 | 0.569 |
| tAlaAGC2-11 | 426 | 16 | 11 | 3.005 | 0.569 |
| tAlaUGC1-1* | 770 | 16 | 5 | 0.44 | 0.259 |
| tAlaUGC2-1* | 770 | 16 | 5 | 0.44 | 0.259 |
| tLeuUAA4-1* | 753 | 21 | 7 | 0.601 | 0.362 |
| tLeuCAG1-3* | 639 | 21 | 0 | 0.014 | 0.118 |

<sup>1</sup> Asterisks denote genes that could not be amplified from *Drosophila* genomic DNA.
<sup>2</sup> ATACSeq describes values obtained from (Huynh et al., 2023) that describe the frequency of sequencing reads gathered from the associated tRNA gene’s genomic locus, which act as a measure of chromatin accessibility.
<sup>3</sup> Isoacceptor and isodecoder counts act as indications tRNAs competition; isoacceptors are tRNAs charged with the same amino acid, and isodecoders are tRNAs carrying the same anticodon (P. P. Chan & Lowe, 2008).
<sup>4</sup> Cognate RSCU (relative synonymous codon usage) describes biases toward codon presence within a set of highly expressed mRNAs (P. M. Sharp et al., 1986).
<sup>5</sup> Codon optimality is a tRNA adaptation index metric (tAi), which represents codon optimality, or how well a codon is translated, based on cognate tRNA presence, codon use, and biochemical codon-anticodon interaction properties (Anwar et al., 2023).
<sup>6</sup> tRNA<sup>Ser</sup> mutant toxicity are metrics based on the data presented in Zimmerman et al., 2018 that assessed cells expressing mutant tRNA mistranslation of Ser at all codons to assess mistranslating cells’ ability to compete with non-mistranslating cells within a single population.

### DNA cloning

*Drosophila melanogaster* tSerUGA1-1, tSerAGA2-4, tAlaAGC2-1, and tAlaAGC2-11 were PCR amplified from genomic DNA. All primer sequences can be found in Table S3.

Amplicons contained roughly 300-500 base pairs on both sides of the tRNA gene. Gel purified amplicons were ligated into pGEM T-easy cloning vector and assessed via Sanger sequencing. Several tRNA genes were unable to be PCR amplified from genomic DNA: tSerAGA1-1, tSerGCU2-2, tSerGCU2-4, tAlaTGC2-1, tLeuTAA4-1, and tLeuTAG1-2.

T-easy or IDT plasmids containing tRNA genes were used as templates for two-step mutagenesis PCRs to generate mistranslating tRNA mutants. Mutagen amplicons were ligated into T-easy cloning vectors and confirmed via Sanger sequencing. Generation of mistranslating mutant tRNAs with either G9A or G26A secondary mutations was accomplished via a second round of two step mutagenesis PCR reactions using mistranslating mutant tRNAs as a template.

Amplicons of secondary mutation generating PCRs were again ligated into T-easy vector for subsequent cloning into both *Drosophila* and yeast expression vectors. Additional mistranslating tRNA mutants created exclusively for assessment in yeast were mutagenized via Q5 mutagenesis PCR using inducible yeast tRNA expression plasmids as templates and confirmed via Sanger sequencing.

tRNA mutant T-easy vectors were restriction enzyme digested with either NotI or EcoRI to be ligated into *Drosophila* or yeast expression vectors, respectively. The *D. melanogaster* expression plasmid pUC-IDT-pattB was generated previously (Isaacson et al., 2024) and contains an attB recombinase site, a mini-white rescue gene for screening purposes, as well as a FRT-site flanked multiple cloning site to allow flippase dependent removal of the transgenic tRNA. The yeast expression plasmid YcPlac33 (Ura) was a generous gift from Dr. Chris Brandl. Neither *Drosophila* nor yeast expression plasmids contain promoters or terminators.

To generate inducible tRNA expression plasmids for yeast, the doxycycline / tetracycline inducible tRNA expression system (Berg et al., 2021) was utilized. tRNA mutants were PCR cloned 5’ HindIII and 3’NotI into the plasmid 4700 gifted by Dr. Chris Brandl. This plasmid is derived from Ycplac111 (Leu) and contains a CYC1 promoter whose activity is induced by the presence of tetracycline. Constitutively expressed tetracycline from the yeast strain TetR (gifted by Dr. Brandl) allows for constant suppression of transgenic tRNA expression. Addition of doxycycline to growth media causes the release of tetracycline from the CYC1 promoter, resulting in inactivation of the promoter and uninhibited expression of the adjacent tRNA.

### Yeast strains and transformation

Yeast Y33-tRNA plasmids were transformed into the strain BY4742 using conventional lithium acetate transformation and plated on SD plates lacking uracil. Yeast inducible-tRNA expression plasmids Y11-CYC1-tRNAs were transformed into the TetR yeast strain (a derivative of BY4742 with genomically integrated constitutively expressing tetracycline and uracil transgenes) using lithium acetate transformation and plated on SD plates lacking uracil and leucine.

### Yeast growth assay

Yeast strains were grown in quadruplicate overnight at 30°C with the appropriate SD media and 250RPM orbital shaking. Culture density was measured by diluting each culture 15µL in 285µL water in a 96 well clear cell culture plate and measuring OD^600^ with a Tecan Infinite M200 plate reader. Diluted culture OD was used to calculate ODs of overnight cultures. 300µL of the appropriate SD selection media was placed into each well of a 96-well clear cell culture plate. Calculated volumes of yeast strain overnight cultures were added to 300µL selective media in a 96 well plate to obtain a starting OD^600^ of 0.1. Depending on the strains used, selective media contained 1.0ug/mL of doxycycline to induce tRNA expression. Following OD normalization in new media, 96 well plates were transferred to a BioTek Epoch 2 microplate spectrophotometer programmed to read OD^600^ at 15-minute intervals for 24 hours, with agitation and temperature set to 30°C.

OD^600^ data output from the plate reader was exported and used with the R package Growth Curver to calculate both doubling time and area under curve as proxies for cell health. Relative growth (either doubling time or area under curve) was determined using the growth of an individual strain relative to the average growth for the appropriate control strain. Relative growth metrics were statistically analyzed in Prism software with a two tailed ANOVA followed by pairwise Dunnett’s multiple comparisons tests accounting for multiple comparisons.

### Mass spectrometry

Yeast cells containing inducibly-expressed mistranslating tRNA variants were inoculated in the 3mL selected media lacking doxycycline overnight. OD^600^ was measured for overnight cultures, and a calculated volume of each culture was added to 7mL of selective media such that OD was 0.1. Importantly, selective media contained 1ug/mL doxycycline to allow transgenic tRNA expression. Normalized OD cultures in new doxycycline-containing media were grown at 30C with 250RPM orbital shaking for 24 hours. Cultures were then pelleted, washed with ddH2O, and flash frozen in liquid nitrogen.

Sample protein isolation, preparation and liquid chromatography tandem mass spectrometry were performed by Bioinformatics Solutions Inc. Detailed procedures can be found in supplemental information. Briefly, yeast cells were lysed via sonication and quantified via BCA assay, after which 10µg of protein per sample was digested with trypsin. Peptides were analyzed by nanoflow LC-MS/MS using an EASY-nLC 1000 (ThermoFisher, Massachusetts, USA) coupled to a timsTOF Pro (Bruker Daltronics, Bremen, Germany) equipped with a CaptiveSpray source (Bruker Daltronics, Bremen, Germany). Reverse-phase chromatographic separation was performed at 300µL/min using a 15cm reversed-phased column with a 75 µm inner diameter filled with Reprosil C18 (PepSEP, Bruker, Germany). The mass spectrometer was operated across a 100-1700 m/z mass range and ion mobility range of 0.85-1.30 s/cm2, utilizing collision energy ramping and active exclusion. Data was compiled and analyzed using PEAKS software. Mutant peptides were only included in analysis if the particular substitution occurred greater than 3 amino acids distance from the edge of a peptide’s cleaved ends, such that no ambiguous identification could occur. The number of mistranslated peptides are reported as the number of unique peptide species identified containing the expected amino acid substitution that could be confidently distinguished from their wildtype counterparts. Frequency of mistranslation is calculated as an average of each mutant peptide’s relative abundance compared to its wildtype peptide counterpart.

### Drosophila melanogaster mistranslating mutant tRNA model creation

*Drosophila* expression plasmids were concentrated by growing transformed Eschericia coli strains in 30mL cultures of LB amp overnight. Cultures were aliquoted into smaller volumes and plasmids harvested with NEB DNA mini prep kits. Mini prep procedure was standard with the exception of DNA binding to column matrices - all aliquots of the same culture were passed through the same matrix to highly concentrate it with identical plasmid. From here, matrices were eluted with ddH20 and assessed for concentration. Each plasmid was normalized to a concentration of 700ng/μlL and a ⅙ volume of sterilized blue food dye was added.

The *Drosophila* strain *y^1^ M{RFP3xP3.PB GFPE.3xP3=vas-int.Dm}ZH-2A w*; PBac{y+- attP-3B}VK00037* (Bloomington *Drosophila* Stock Center stock #24872) was used to generate tRNA mutant transgenics, chosen for its intergenic att3B/VK00037 landing site, *white* eye pigment mutation, and X-linked germline tissue expression of the phi-C31 integrase required for integration of genetic material from the plasmid into the att3B landing site. Embryos of strain #24872 were harvested using an egg collection apparatus. Following a 10-20 minute laying period, embryos were collected from apple juice plates using fine paint brushes and aligned along a glass cover slip. Embryos were then air-dried momentarily to allow fixation to the cover slip and subsequently submerged in neutral oil to prevent desiccation. Injection of the tRNA expression plasmids into embryos was accomplished with the use of ultra-fine needles and a micro injector. Injection of all embryos on a cover slip occurred within 20 minutes to prevent excessive development. Cover slips were then inserted into a standard *Drosophila* food vial such that the anterior ends of the embryos were adjacent to the top of the food.

Following injection, individuals that survived into adulthood were collected and crossed to one additional injection survivor of the opposite sex. Progeny of these single parent crosses were screened for eye colour - those that had integrated the injected plasmid display yellow eyes, while those that do not contain the transgenic retain the white eye colour expected of stock #24872. Transgenic survivors were then crossed to virgins of the genotype *w-;CyO/Sco* such that stable stocks could be created. Presence of transgenic tRNA was confirmed by harvesting DNA, PCR amplifying an amplicon specific to the integrated plasmid, and Sanger sequencing the amplicon.

### Drosophila viability assay

For each transgenic tRNA variant, 5 males of the genotype *w*-;tRNA/*CyO* were crossed to 20 virgin females of wildtype Canton-S (CS) strain for a single biological replicate. Seven biological replicates were conducted per transgenic tRNA. Parent flies were permitted to lay eggs for seven days before being transferred to a new food vial. F1 progeny were genotyped, sexed, and counted every two days for 10 days to prevent inclusion of F2 individuals.

Proportions of individuals inheriting the transgenic tRNA were compared to those inheriting the *CyO* balancer chromosome. Deviations from proportions expected of mendelian ratios was used as a proxy for developmental deficiencies during larval and pupal stages. Such mendelian deviations were calculated relative to the SerUGA1-1 wildtype tRNA strain.

### Drosophila stock maintenance

All stocks, either created here or obtained elsewhere, were reared on standard Bloomington *Drosophila* food media, housed within incubators at 24°C, 70% humidity, and a 14:10 light:dark cycle.

## Results

### Impact of different forms of mistranslation on yeast growth

To assess which characteristics of mistranslation are important in eliciting biological effects, various mistranslating anticodon mutants of the *Drosophila* tSerUGA 1-1 gene were integrated into yeast cells where growth rate was measured as a proxy for cell health. This initial screen included 7 anticodon mutants of the tSerUGA1-1 gene that mistranslate either Gly, Ile, Leu, Pro, Thr, Trp, or Val codons, while yeast expressing wildtype copy of said tRNA^Ser^ gene served as a control. Of the chosen anticodon mutants, Ile-AAT, Leu-TAA, and Trp-CCA produced statistically significant reductions in growth rate when expressed constitutively in yeast cells (Figure 1, top). The average growth rates of other groups trend toward similar reductions but display substantial variation, owing to a subset of individual replicates displaying impaired growth, while others appear unaffected.

Given the inviability of specific mistranslating variants in previous models (Berg et al., 2017), as well as the intragroup variation observed here, concerns arose over the occasional emergence of suppressor mutations that ameliorate the toxic effects of the mistranslating tRNA. Such phenomena could also contribute to the lack of growth deficit observed in other groups - strong mistranslation could hypothetically incite suppressor mutation emergence in each replicate. To address these concerns, a number of approaches were employed. First, we adopted the doxycycline inducible tRNA expression system (Berg et al., 2021). In this system, mutant mistranslating tRNAs are cloned immediately 3’ to the TetO-CYC1 promoter, whose activity competitively inhibits tRNA transcription machinery. The activity of the TetO-CYC1 promoter is dependent on the presence of tetracycline, which is constitutively expressed by a genomically integrated transgene in the yeast strain BY4742. At the start of the assay, the growth media is supplemented with doxycycline, which binds tetracycline and frees the TetO-CYC1 promoter, halting its competitive inhibition and therefore permitting transcription of the adjacent tRNA. Use of this system allows for temporal control of mistranslation, limiting its occurrence to only the assay (24 hours) as opposed to the entirety of the transformation and liquid culture phases (3-5 days). This constrained timescale greatly limits the possibility of suppressor emergence.

Use of the doxycycline inducible tRNA expression system to limit mutant tRNA to the growth assay itself produces results distinct from constitutive expression of the same mutants. Each of the assessed mutants display impaired growth relative to the wildtype (WT) tRNA control (Figure 1, bottom), as well as considerably reduced intra-group variation. Although all assessed mutant tRNAs confer growth impairment relative to the WT tRNA control, the magnitude of growth arrest differs substantially between mutants. Pro-UGG and Thr-AGT anticodons produce the least change to growth rate at roughly 90-95% of WT control, while Gly- ACC, Ile-AAT, and Val-AAC all reduce growth to roughly half of the WT control. Growth rate change of Leu-TAA mutants is of an intermediate severity between these two groups, growing at roughly 75% of wild type control.

### Impact of mistranslation frequency on yeast growth

Next, to assess the effects of mistranslation frequency on growth outcomes, we generated each mistranslating tRNA variant in two additional conditions that tailor the amount of mistranslation occurring. The inclusion of secondary mutations that destabilize the mutant tRNA’s structure ultimately reduces said tRNA’s participation in translation, thus lowering the amount of mistranslation that occurs. The three experimental conditions affecting frequency of mistranslation are as follows: a mutant mistranslating tRNA with no secondary mutation (most frequent mistranslation), a mutant mistranslating tRNA with a G26A secondary mutation (moderately frequent mistranslation), or a mutant mistranslating tRNA with a G9A secondary mutation (least frequent mistranslation)(Berg et al., 2017). These mistranslation frequency conditions were generated for both constitutive long-term and induced short-term expression conditions. For anticodon mutants that caused growth deficits when no secondary mutation was present, the addition of secondary mutations reduced said growth deficit to wildtype-like levels in the case of G9A mutants, and slightly below wildtype for G26A mutants (Figure 2). Anticodon mutants without secondary mutations that did not produce growth deficits were largely unchanged by the addition of secondary mutations.

**Figure 2.**
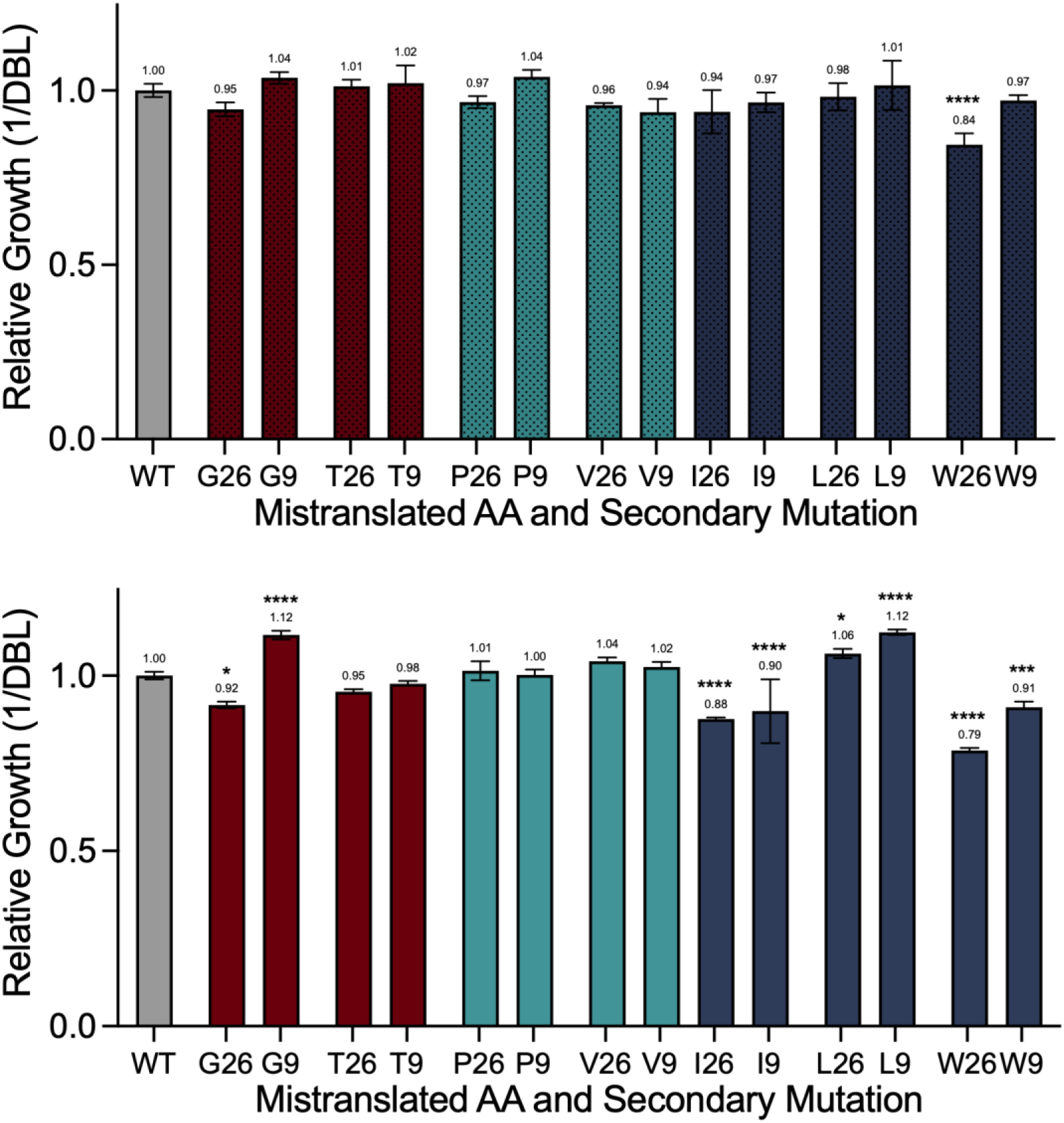
Growth deficits imbued by different forms of mistranslation are ameliorated by the addition of secondary mutations. Forms of mistranslation are grouped by colour according to qualitative magnitude of biochemical difference to serine: Gly (G) and Thr (T) are a low magnitude of change (maroon), Pro(P) and Val (V) comprise a medium magnitude of change (teal), whereas Ile (I), Leu (L), and Trp (T) are a high magnitude of change (navy). Variants containing the G26A mutation mistranslate at a moderate frequency (denoted by x-axis “26” label), while those containing a G9A mutation mistranslate at a lower frequency (denoted by x-axis “9” label). Top: constitutive expression of tRNA variants. Bottom: expression of the same tRNA variants limited exclusively to the 24-hour growth assay via inducible tRNA expression system. Group means are listed as annotations above each bar (n=4). Error bars denote standard deviation. Statistical significance as determined by one-way ANOVA and subsequent post-hoc pairwise Dunnet’s multiple comparisons tests is marked by asterisks (**** denotes p<0.00001, *** denotes p< 0.0001, ** denotes p<0.01, * denotes p<0.05).

These results largely hold true when secondary mutant, low-frequency mistranslation, is restricted to the shorter 24 hour assay period via the inducible tRNA expression system. Assay- only expression of tRNAs with secondary mutations results in substantial magnitude decreases of growth deficits compared to tRNAs with the same mistranslating anticodon and no secondary mutation (Figure 2, bottom). Many instances of low-frequency mistranslation in this short-term expression context demonstrate a mild increase in growth rate when compared to wildtype control. Although there are statistically significant reductions of growth rate compared to WT for several mild-frequency mistranslating tRNA variants expressed in the short-term, these differences are of smaller magnitude than their higher-frequency counterparts (0.84 relative growth for moderate mistranslation of Trp via G26A mutation, compared to 0.34 relative growth for the same form of mistranslation at a higher frequency; Figure 1).

The results above demonstrate an important effect for the identities of mistranslated amino acids in eliciting growth deficits. However, nearly all amino acids are coded by multiple synonymous codons that differ in both frequency of use and presence within specific protein products. To assess whether specific codon identity affects the outcomes of mistranslation, and to begin inferences as to how these hypothetical differences may arise, we created an additional group of tRNA mutants that result in the same amino acid substitutions assessed above but mistranslate alternative synonymous codons (see Table 1). As tryptophan is only coded by one codon/anticodon, it was excluded from these experiments.

For several of the tRNA mutant groups decoding the same amino acid, variation in growth deficit severity is observed among the alternative anticodon mutants (Figure 3). Of the novel anticodon mutants assessed here, Gly-TCC and Leu-GAG dramatically decrease the relative growth deficit displayed by their synonymous codon peers. Alternate Pro and Thr anticodons are consistent with their synonymous codons assessed previously.

**Figure 3.**
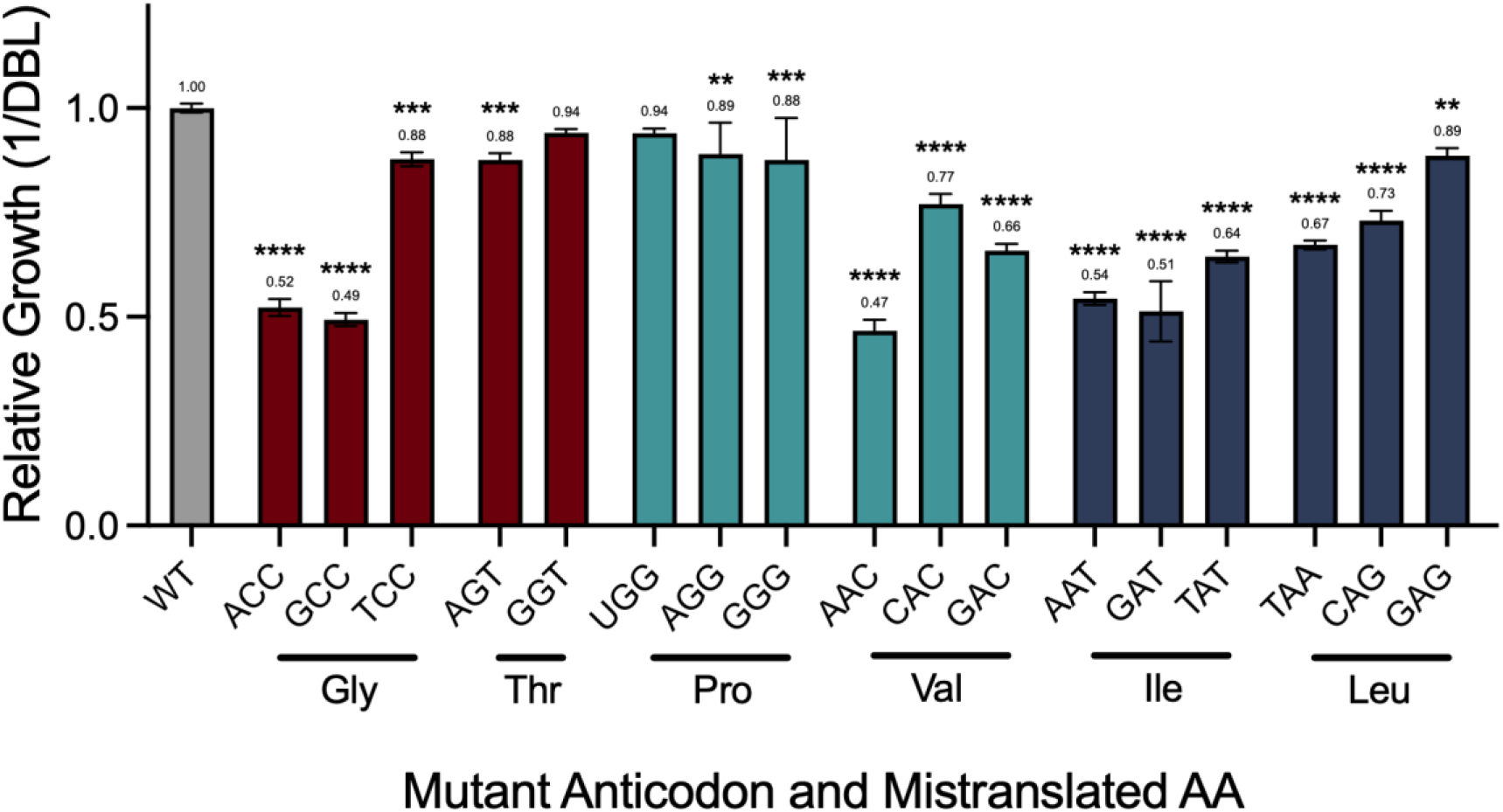
Mistranslation of synonymous codons results in different growth outcomes. Doubling time growth of cells expressing each mutant is reported relative to cells expressing a wildtype copy of the same tRNA gene. Expression of mistranslating mutant tRNAs is limited to the 24 hour growth assay via the doxycycline inducible tRNA expression. Forms of mistranslation are grouped by colour according to qualitative magnitude of biochemical difference to serine: Gly and Thr are a low magnitude of change (maroon), Pro and Val comprise a medium magnitude of change (teal), whereas Ile, Leu, and Trp are a high magnitude of change (navy). Group means are listed as annotations above each bar (n=4). Error bars denote standard deviation. Statistical significance as determined by one-way ANOVA and subsequent post-hoc pairwise Dunnet’s multiple comparisons tests is marked by asterisks (**** denotes p<0.00001, *** denotes p< 0.0001, ** denotes p<0.01). Growth curves can be found in Supplementary Figure S2.

While we and others have demonstrated a breadth of phenotypes elicited from one tRNA gene bearing many different mutations, and thus different forms of mistranslation, comparatively little is known regarding the influence on such phenotypes imbued by the specific tRNA gene bearing the mistranslating mutation. Here, we attempt to further our understanding of how different isoacceptor tRNA genes may influence phenotypes when harboring the same mutation. Again, the data presented above were generated using mutants of the *D. melanogaster* tSerUGA1-1 gene. An alternative *Drosophila* tRNA^Ser^ gene, tSerAGA2-4, was mutagenized to contain Pro-UGG and Gly-ACC anticodons and was subsequently transformed into yeast strains and assessed for growth phenotypes.

Interestingly, tSerAGA2-4 bearing the Gly-ACC anticodon produces a decrease in growth rate (Figure 4). While this mirrors the phenotype of the tSerUGA1-1 Gly-ACC mutant in Figure 1, there is a marked reduction in the magnitude of growth deficit observed in the tSerAGA2-4 mutant. However, both tRNA^Ser^ genes display no growth effects with the presence of the Pro- UGG anticodon.

**Figure 4.**
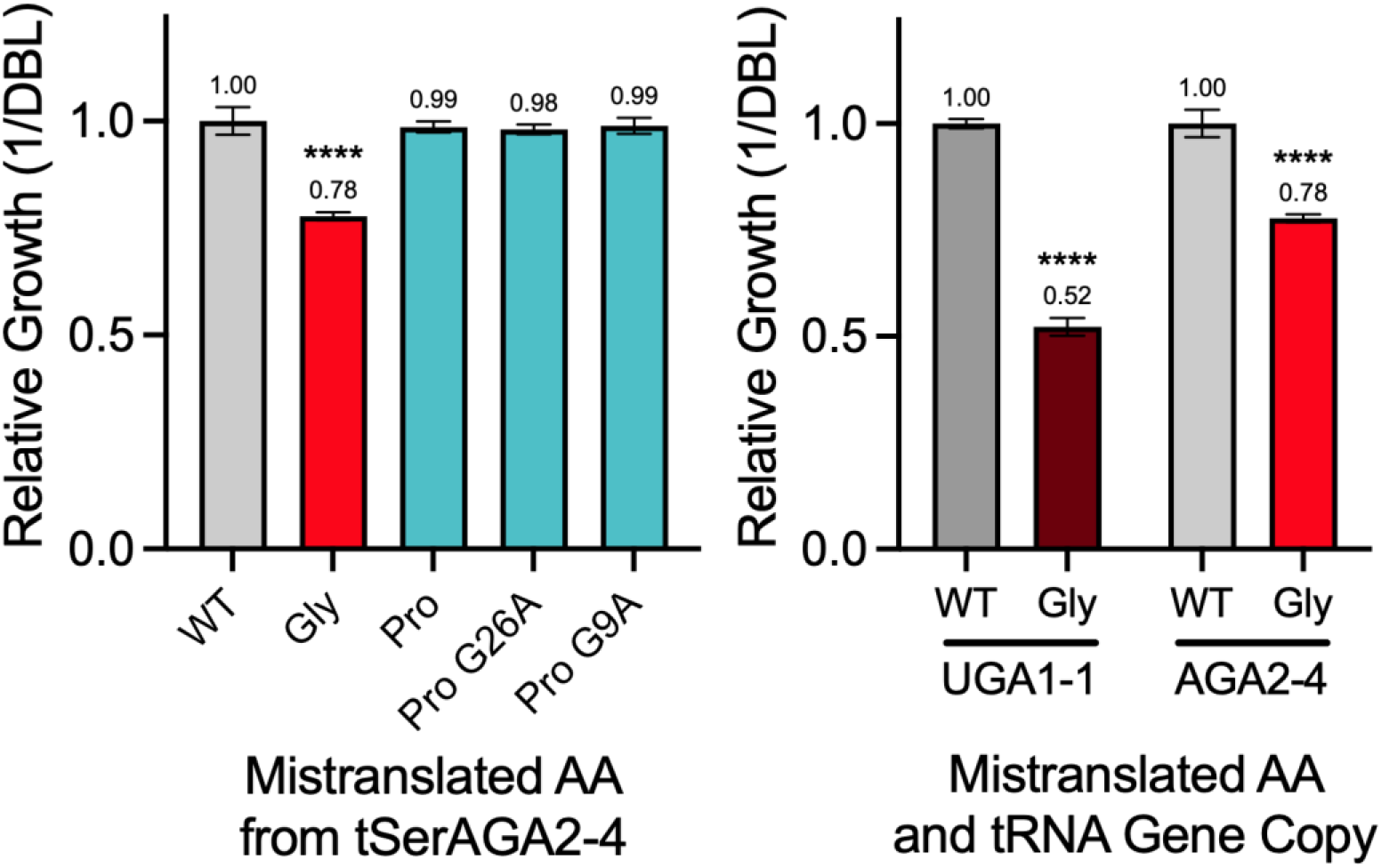
Alternate copies of tRNA^Ser^ genes bearing the same mistranslating mutations confer different magnitudes of growth deficits. Left: doubling time growth of tRNA variants of the tSerAGA2-4 *Drosophila* tRNA gene relative to cells expressing a wildtype copy of said tRNA gene. Presence of secondary mutations that reduce mistranslation frequency are noted in x-axis labels. Right: doubling time growth of cells expressing two distinct tRNASer copies bearing identical mistranslating anticodon mutations relative to their corresponding wt tRNASer gene. Gene copies are grouped along the x-axis and denoted by darker (tSerUGA1-1) and lighter (tSerAGA2-4) colour schemes. Group means are listed as annotations above each bar (n=4). Error bars denote standard deviation. Statistical significance as determined by one-way ANOVA and subsequent post-hoc pairwise Dunnet’s multiple comparisons tests is marked by asterisks (**** denotes p<0.00001, *** denotes p< 0.0001).

Of the forms of mistranslation assessed above, all are similar in their use of serine as the mistranslated amino acid replacing the coded residues. Other non-serine forms of mistranslation are possible and near certainly different in their array of resulting phenotypes. Of note, tRNA^Ala^ and tRNA^Leu^ are similar to tRNA^Ser^ in that their anticodon sequences are not identity elements used for aminoacylation, and thus mutations to the anticodons of these genes also produces mistranslation. With the eventual goal of integrating tRNA^Ala^ and tRNA^Leu^ mistranslating variants into *D. melanogaster* models of mistranslation, we begin by compiling a list of tRNA genes to mutagenize and assess in yeast as a high throughput model. The genes chosen are also outlined in Table 2, and were chosen largely for their chromatin accessibility as a proxy for anticipated high expression (Merrill et al., 2022; Milon et al., 2014). Unfortunately, several of the chosen genes were unable to be amplified from genomic DNA, and thus planned experiments proceeded with a smaller subset of tRNA^Ala^ genes.

Of the viable tRNA gene candidates, tAlaAGC2-1 was mutagenized to encode several anticodons to begin assessment of the effects of different forms of Ala mistranslation: Thr-AGT, Thr-GGT, Pro-UGG, and Val-AAC. Subsequently, the anticodon of an additional tRNA^Ala^ gene, tAlaAGC2-11, was also mutagenized to Val-AAC. All of the mistranslating mutations in either of the examined tRNA^Ala^ genes produced no growth deficits when expressed in *Saccharomyces cerevisiae* (Figure 5).

**Figure 5.**
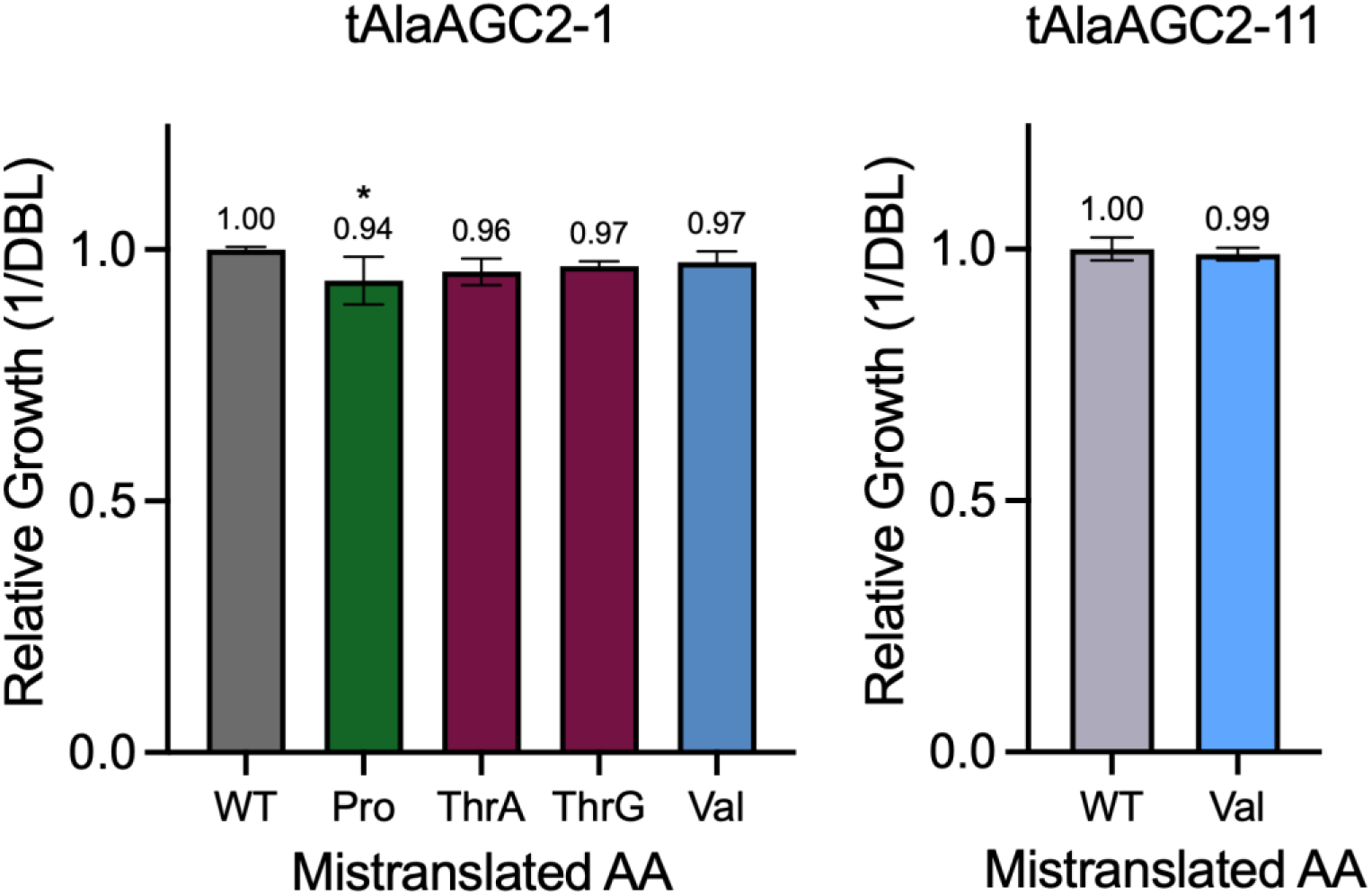
Mutants of *Drosophila* tRNA^Ala^ genes confer no growth change when expressed in yeast cells. Left: doubling time growth of yeast cells expressing mutants of the *D. melanogaster* tAlaAGC2-1 gene relative to cells expressing a wildtype copy of the same tRNAAla gene. All variants were expressed exclusively during 24 hour growth assay via the doxycycline inducible tRNA expression system. Anticodon mutants correspond to Pro-UGG, Thr-AGT, Thr-GGT, and Val-AAC. Right: doubling time growth of yeast cells expressing mutants of the *D. melanogaster* tAlaAGC2-11 gene, relative to cells expressing a wildtype copy of said gene. Group means are listed as annotations above each bar (n=4). Error bars denote standard deviation. Statistical significance as determined by one-way ANOVA and subsequent post-hoc pairwise Dunnet’s multiple comparisons tests are marked by asterisks (**** denotes p<0.00001, *** denotes p< 0.0001).

### Confirmation and quantification of mistranslation

In order to confirm the growth phenotypes observed thus far can confidently be attributed to anticodon mutation-bearing tRNAs expressed in these cells resulting in mistranslation, we assessed mutant peptide presence via mass spectrometry. Whole protein lysate collected from cells expressing either Gly-AAC, Val-ACC, Ile-AAT, and a WT control tRNA were used to provide a range of phenotypes for this analysis. On average, roughly 38,000 peptides were confidently identified. For each tRNA mutant, the number of peptides containing the expected amino acid substitution was assessed, whereas the WT tRNA control group was assessed for each of the amino acid substitutions of interest. The WT control tRNA does not cause any significant increase in mutant peptide presence for any of the assessed substitutions. In contrast, each of the mistranslating tRNA groups significantly increases the peptides containing the expected amino acid substitution (Figure 6). The assessed mistranslating tRNAs also differ in their frequency of mistranslation, measured as the proportion of mutant peptide presence relative to the corresponding wildtype peptide. Mistranslation frequency is highest among the Val-AAC mutant, while Gly-AAC and Ile-AAT are slightly lower.

**Figure 6.**
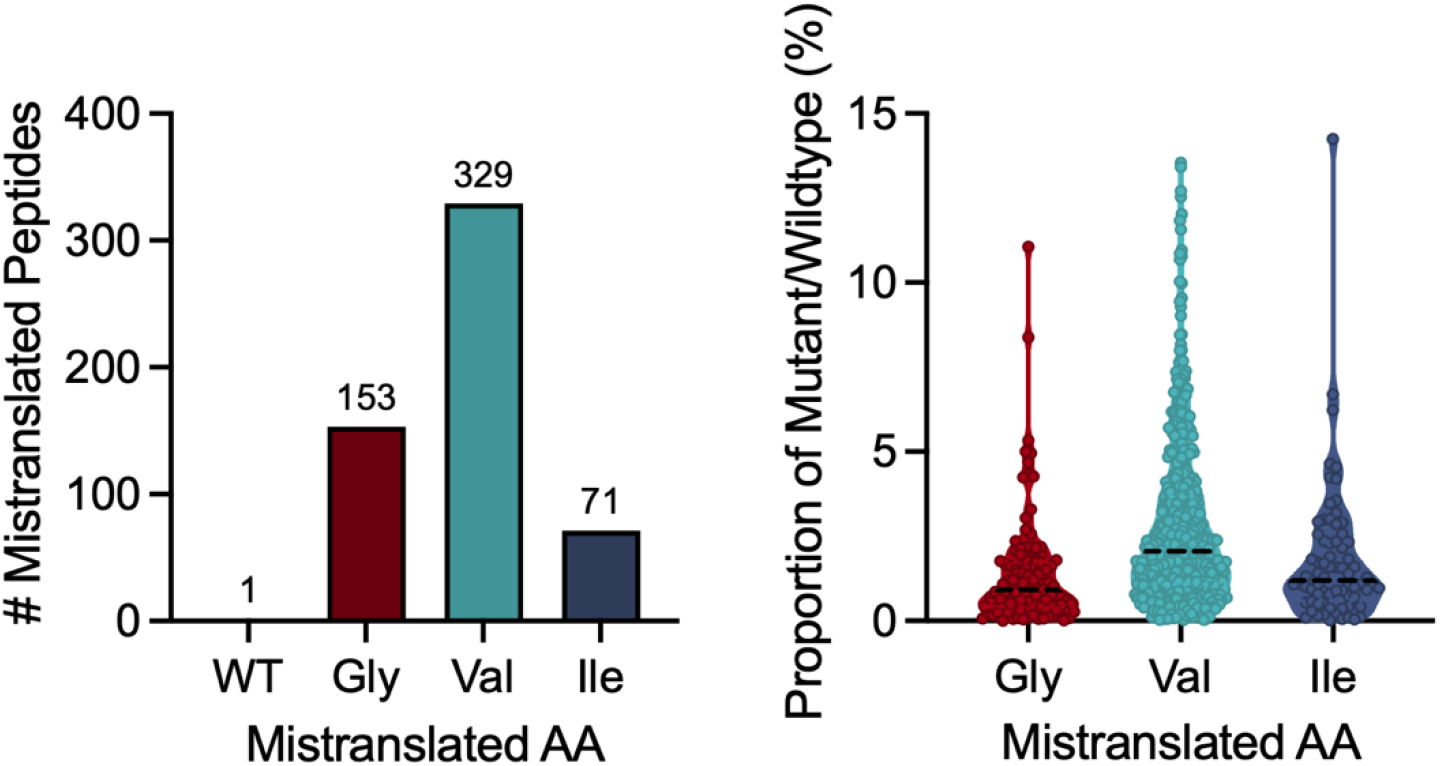
Yeast cells expressing mistranslating tRNA variants produce increased numbers of mutant peptides. Left: Number of mutant mistranslated peptides identified by mass spectrometry of whole protein lysates collected from yeast cells expressing *Drosophila* tSerUGA1-1 variants. WT tRNA genotype was assessed for all three forms of mistranslation, whereas each mistranslating mutant was only assessed for mutant peptides corresponding to the anticipated substitutions. Right: mistranslation frequency as measured by the proportion of mutant peptides relative to wildtype isoforms. Each dot represents an individual mutant/wildtype peptide ratio, while the thick dashed line represents mean mistranslation frequency and the dotted line represents one standard deviation from the mean.

### Translation of novel mistranslating tRNA variants into Drosophila melanogaster

Previously, we designed tRNA^Ser^ mistranslating tRNA mutants Val-AAC and Thr-AGT, both with the G26A secondary mutation, which were integrated into *D. melanogaster* to assess the impacts of mistranslation in a multicellular model (Isaacson et al., 2024). Several other tRNA variants bearing G26A mutations were also integrated into *D. melanogaster* but were abandoned due to the inability of mass spectrometry to detect elevated mutant peptides, despite some displays of developmental phenotypes.

Here, we attempt to integrate additional mistranslating tRNA variants with prospective differences in mistranslation frequency to those established previously, namely mutants without secondary mutation and those with a G9A mutation allowing the mildest amount of translational participation. The list of mutant tRNAs injected, number of embryos injected, survivor counts, and the number of transgenic individuals produced are all outlined in Table 3. Unfortunately, several of the attempted mutant mistranslating tRNAs were unable to be established as transgenic lines in *Drosophila*. In particular, mistranslating tRNAs without secondary mutation produced virtually no transgenic individuals despite thousands of injected embryos. A smaller number of mutant tRNAs bearing the G9A secondary mutation, Leu-TAA and Thr-AGT, were also unsuccessful in their integration into *Drosophila*. Pro-UGG mutants were not attempted, as they constitute a previously established model (Isaacson et al., 2022).

**Table 3.** Some forms of mistranslation appear inviable in *Drosophila melanogaster*.

| tRNA<br>Variant | #<br>Injected | Male<br>Survivors | Female<br>Survivors | Total<br>Survivors | # With<br>Transge |
|---|---|---|---|---|---|
| Gly | 681 | 22 | 17 | 39 | 0 |
| Ile | 687 | 17 | 21 | 38 | 0 |
| Thr | 937 | 23 | 29 | 52 | 0 |
| Trp | 719 | 25 | 17 | 42 | 0 |
| Val | 832 | 30 | 24 | 54 | 0 |
| Gly G9A | 531 | 28 | 19 | 47 | 3 |
| Ile G9A | 784 | 46 | 38 | 84 | 2 |
| Leu G9A | 540 | 16 | 21 | 37 | 0 |
| Thr G9A | 546 | 16 | 17 | 33 | 0 |
| Trp G9A | 1194 | 20 | 15 | 35 | 2 |

Of the several SerUGA1-1 mutant tRNAs attempted, a select few were successfully integrated into the *D. melanogaster* genome: Leu-CAA, Gly-ACC with G9A, Ile-AAT with G9A, Trp-CCA with G9A, Val-AAC with G9A, and a wildtype tSerUGA1-1 control. Despite the successful establishment of a Gly-ACC with G9A transgenic line, the line proved short lived, dying shortly after establishment and thus was excluded from viability assessment. Given that previous *Drosophila* models of mistranslation regularly displayed developmental lethality, we assessed developmental viability of each of the newly established transgenic tRNA lines. This was accomplished using heterozygous mutant tRNA stocks wherein the homologous chromosome contained a visible marker, and crossing these stocks to a wildtype strain. Progeny of these crosses were genotyped, tallied, and compared to both expected mendelian ratios as well as the ratios of genotypes observed for the wildtype tSerUGA1-1 crosses, while also considering sex and time of eclosion as important co-variates. Perhaps surprisingly, viability of mutant tRNA stocks display no statistical difference to that of the wildtype tRNA strain (Figure 7). These consistencies between groups are true of the raw count data as well as group proportions modeled with mixed linear regression accounting for sex and time to eclosion.

**Figure 7.**
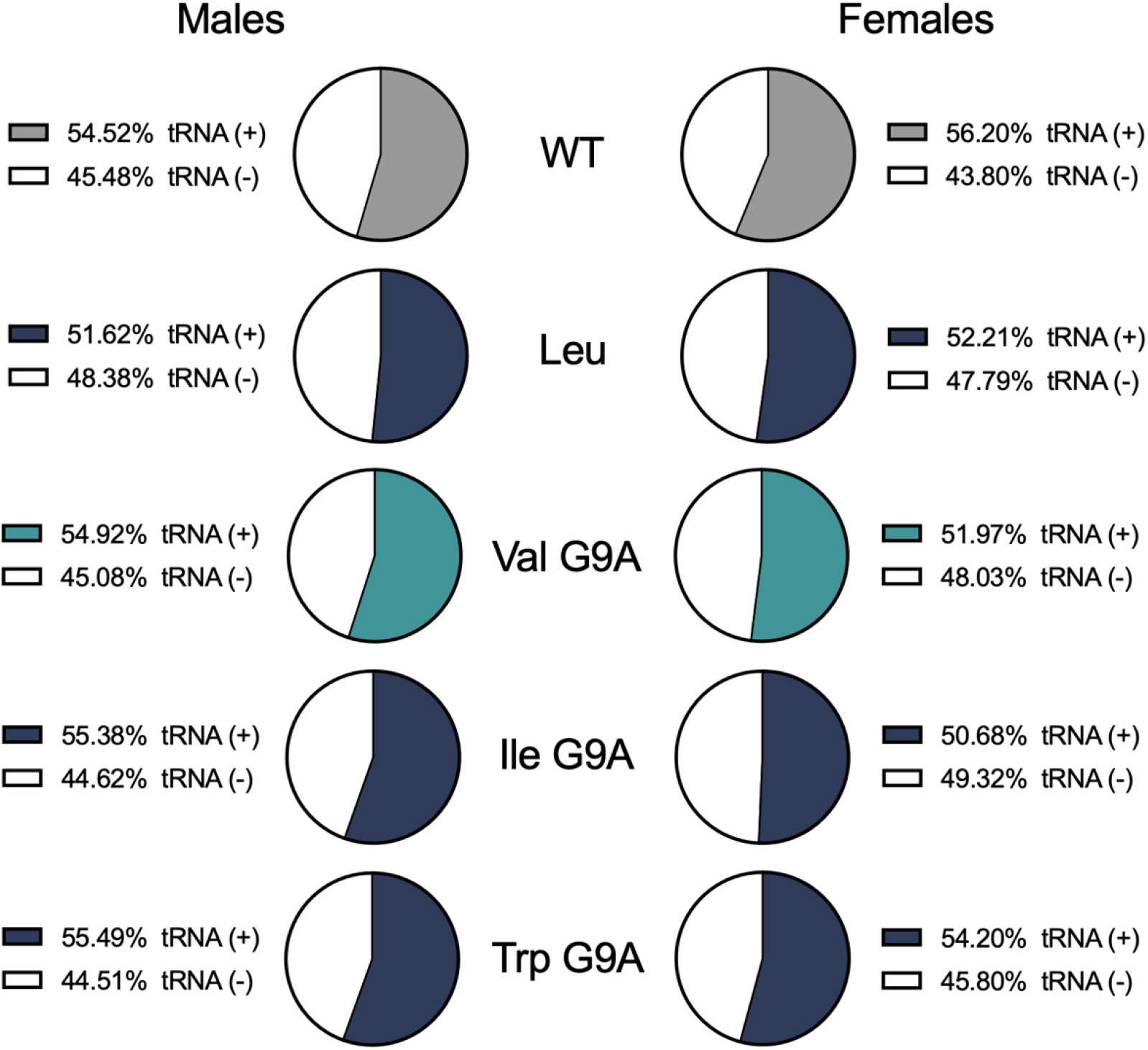
Different forms of mistranslation have no effect on survival to adulthood in *Drosophila melanogaster*. Each pie chart represents the proportion of individuals that survive to adulthood that contain mutant tRNAs (colour fill) and individuals that do not (white fill). Pie charts on the left represent male adults, while those on the right represent female adults. Mistranslating tRNAs are grouped in rows. For each mistranslating tRNA group, deviations from the proportions observed in the wildtype tRNA control group would constitute differences in developmental viability. Statistical analyses via pairwise chi-square or mixed linear regression with eclosion time as a random variable revealed no such deviations from WT genotype proportions.

## Discussion

### Yeast model of Drosophila tRNA mutant mistranslation

Here, we assess how the effects of mistranslation are dictated by various characteristics of the specific mistranslating mutation and tRNA gene identity, referred to here collectively as the form of mistranslation. Based on this data, as well as similar experiments by others, no single characteristic of mistranslating mutations can be identified as most important in determining a mistranslating mutation’s deleteriousness (Berg et al., 2021; Davey-Young et al., 2024; McDonald et al., 2025; Zimmerman et al., 2018). However, general rules and parallel prospective hypotheses can be gleaned from these works.

The severity of biochemical difference between the substituted amino acids appears to be a reasonable predictor of mistranslating tRNA mutant toxicity. The most severe biochemical changes to Ser, namely from Trp, Leu, and Ile, all produce substantial growth deficits in both induced and constitutive expression contexts. Moderate and milder biochemical differences are more complicated to interpret given the variety of growth changes stemming from amino acid changes similar in severity. Of the more moderate biochemical changes, Val and Pro differ substantially in their effects. Val produces significant growth arrest when inducibly expressed, but no statistically significant difference when constitutively expressed. Contrastingly, mistranslation of Pro produces no growth change in either expression context. A similar lack of consistency is true of the mildest biochemical changes - mistranslation of Thr produces no growth deficits, while Gly is highly toxic when inducibly expressed.

In addition to growth deficits, biochemical differences in substituted amino acids seem to also influence the probability for suppressors of mistranslation to emerge. Of the forms of mistranslation toxic when inducibly expressed, the same mutant expressed constitutively often displays exceptionally high variation between replicates (ex. Gly, Leu, and Trp; Figure 1). The most plausible explanation for the high replicate variation is the emergence of mistranslation suppressors in some but not all individual cultures, or a spectrum of suppression among each.

This high variation could cause many of the forms of mistranslation to fail to reach statistically significant differences from wildtype. Interestingly, the most severe biochemical changes from serine (Trp, Leu, and Ile), maintain their statistically significant difference to wild type when constitutively expressed, despite there being more severe deficits from other mutants when inducibly expressed. This may indicate that these forms of mistranslation are less able to become suppressed. Moderate biochemical changes such as Val and Gly, although highly toxic when inducibly expressed, are occasionally capable of growing at near wildtype rates when constitutively expressed, implying a potential propensity for suppression.

It is also clear that biochemical differences between mistranslated amino acids alone is insufficient to explain the breadth of phenotypes observed. For amino acid changes that occasionally result in toxicity, not all anticodons belonging to the corresponding isoacceptor group are equal in their potential for negative outcomes. Within isoacceptor groups, it appears that the more toxic anticodons are often those that are most frequently used, as indicated by relative synonymous codon usage or tAi codon optimality metrics (Table 1, Table S2)(Anwar et al., 2023; Berg et al., 2021). This stands to reason, as more frequent codon usage should coincide with more mistranslated protein products and their downstream effects. More difficult to explain are the occasional exceptions to this pattern, wherein mistranslation of specific rare and non- optimal codons is toxic, while mistranslation of other similarly rare codons is tolerated well. The distinguishing factor between these scenarios may be the specific proteins in which mistranslation is occurring. Previous literature has demonstrated that the use of rare and non- optimal codons can be functionally adaptive by slowing translational speed to allow co- translational folding (Bae & Coller, 2022; Komar, 2021; Saunders & Deane, 2010; C. H. Yu et al., 2015; Zhou et al., 2013). Mistranslation of the specific rare codons present within these pace- based co-translationally folded protein residues may result in the misfolding of such motifs.

Given that such proteins are also rare, the mistranslation of a fraction of their protein pool may significantly affect their functional capacity. Through such a mechanism, the effects of specific forms of mistranslation may be dictated more by “client protein” identity than general rules regarding amino acid change severity or global mistranslation frequency.

The toxicity of mistranslating tRNA mutants assessed here are largely consistent with similar phenotypes observed in Zimmerman et al., 2018, implying a similar amount of tRNA expression and resulting mistranslation is accomplished by our yeast model compared to theirs. The data presented here and within Zimmerman’s et al in contrast to other experiments, namely Berg et al., 2017, that show significant toxicity for Pro-UGG mutants. This can likely be attributed to differences in expression of the transgenic tRNAs; while Berg’s mutant genes are born of endogenous yeast tRNA genes, those assessed here are trans-species *Drosophila* genes, likely less efficient in yeast transcription, aminoacylation, and/or participation at the ribosome. Further evidence demonstrating Pro-UGG mutants as producing significant biological effects when expressed sufficiently highly can be seen other *Drosophila* models of mistranslation (Isaacson et al., 2022), wherein Pro-UGG mutants with the G9A and G26A secondary mutations cause an array of both negative and positive phenotypes. As such, use of an alternative plasmid that allows increased expression of the transgenic tRNAs is likely to allow for phenotypes distinct from those observed here for the Pro-UGG mistranslating mutants, and potentially also various other forms of mistranslation assessed here.

### Alternative tRNA gene mutants

The effect of mistranslation via two different tRNA^Ser^ tRNAs found that the specific identity of the tRNA bearing the mutation can have drastic effects for the consequences of the mistranslating mutation. Although tSerAGA2-4 is expressed with the same anticodon mutations and from the same genetic context as tSerUGA1-1 it produces a significantly milder growth phenotype. This implies that the two tRNA^Ser^ genes when mutated differ in their relative frequencies of mistranslation. Exactly how this mistranslation frequency difference is manifested requires further investigation. It may be the case that the less severe phenotype that arises from the tSerAGA2-4 is due to lower transcription, potentially due to differences in regulatory sequences that could be present in the ∼300 base pair flanking sequences transgenically included with the tRNAs (Raymond et al., 1985). Alternatively, phenotype effect differences could potentially be caused by the few sequence differences within the tRNA^Ser^ tRNAs themselves.

These sequence differences may alter the transcription factor binding capacity near the A and B box, or the recruitment of transcriptional cofactors. Alternatively, differences in mistranslation frequency may arise from differences in the ability of other factors to act on them post- transcriptionally, such as their corresponding aaRS or tRNA modifying proteins. This may be especially true given the trans-species nature of the tRNA genes assessed here, since there may be a disconnect between specific *Drosophila* tRNA sequences and yeast post-transcriptional modification machinery.

Each of the potential explanations posited here for differences between tRNA^Ser^ gene copies in their mistranslation frequency/growth deficit phenotype can also be applied to the lack of phenotype observed for the tRNA^Ala^ mutants assessed. One may be tempted to interpret these data as evidence that alanine mistranslation is functionally less detrimental than serine mistranslation. However, a wide array of phenotypes were tested for tRNA^Ala^ and tRNA^Leu^ mistranslating mutants, with many demonstrating a significant negative effect of mistranslation (Cozma et al., 2023; Davey-Young et al., 2024). Determining whether the tRNA^Ala^ mutants used here produced no growth phenotypes due to low transcription, irregular modification, or low aminoacylation would require extensive experimentation.

### Detection and quantification of mistranslation

Using a subset of the yeast strains assessing different forms of mistranslation, analysis of mass spectrometry data confirms the occurrence and amount of mistranslation in these models. For instance, this data suggests that Val-AAC mistranslation occurs more frequently than Gly- ACC or Ile-ATT mistranslation, both in the number of mistranslated peptides as well as the relative ratio of mutant peptides to wildtype peptides. Despite this difference in frequency, Val- AAC is not markedly different from either other mistranslating mutant in terms of growth phenotype, further demonstrating frequency of mistranslation is insufficient to explain growth phenotype severity without also considering biochemical differences in mistranslated amino acids or the identity of mistranslated proteins.

Additionally, the number and frequency of mistranslated peptides for some mistranslating mutants differs from what might be expected. Given that Ile is more commonly coded in the genome than both Gly and Val, one might expect mistranslation of ATT, Ile’s most common codon, to result in more frequent mistranslation than Gly and Val mistranslating mutants. This is not the case, and the reason for this discrepancy remains unclear. One potential explanation lies in the corresponding anticodon, AAT, being the most optimal for the isoacceptor group - high optimality equates to a high presence of competitor tRNAs, and thus potentially a lower relative participation at each cognate codon. However, the valine codon chosen for mass spec experiments is also the most optimal and would thus be expected to encounter the same kind of competition. An alternative explanation is the relative presence of each of these codons among highly expressed proteins. Although Ile may be more genomically common, Val and Gly may be more common among proteins that are highly translated, and/or specifically upregulated in response to proteostatic stress. It is worth noting, however, that the identification and quantification of mutant peptides is largely influenced by the ability of the mass spectrometer to detect differences between mutant and the expected wildtype peptides. As such, certain mistranslated peptides may be underrepresented if variants are not confidently distinguishable.

### Drosophila melanogaster mutant mistranslating tRNA model creation

Despite our successes in establishing an array of yeast models of *Drosophila* tRNA gene mutant mistranslation, as well as previous generation of multiple *Drosophila* stocks containing mistranslating tRNA variants, the attempts made here to add to the catalogue of *Drosophila* mistranslation models was largely unsuccessful. However, these efforts do provide valuable insights to be used for future endeavors in creating additional fly models of tRNA variants. Importantly, the tRNA variants assessed here and in Isaacson et al. 2024 were all integrated into the same genomic locus in *Drosophila*. The specific chromatin and/or transcriptional micro- environment provided by this transgenic integration site is nearly certain to affect the transcriptional activity of the inserted mutant tRNAs. As a result, the patterns observed here regarding the viability of various forms of mistranslation are likely to change with the use of alternative integration sites with differing transcriptional activity. The same logic applies to the use of alternative tRNAs whose native transcription may be innately higher or lower than the tSerUGA1-1 used here, thus likely changing the phenotypes they give rise to when mutated to mistranslate.

When considering the tRNA mutants whose integration into *Drosophila* was attempted here, it appears as though those lacking secondary mutations are developmentally inviable, or at least substantially reduced in their viability. A large majority of mutant tRNAs without secondary mutation produced no transgenic individuals following the injection of hundreds of embryos. The single secondary mutation-less transgenic line produced, Leu-TAA, produces no viability phenotypes when compared to its corresponding wildtype tRNA stock. This is in stark contrast to the yeast model for this same mutant, which displays substantial growth deficits (Figure 1). This discrepancy can be interpreted as successful integration of the mutant tRNA, followed closely by the suppression of mistranslation phenotypes. It is also possible that both results are genuine, and that these models differ substantially in their tolerance to this specific form of mistranslation.

Determining which of these two hypotheses is correct would require confirmation and comparison of mistranslation’s continued occurrence and frequency in both models.

Mistranslating tRNA mutants that also contain the G9A secondary mutation were frequently able to produce transgenic *Drosophila*. Of those injected, only Thr-AGT G9A and Leu-TAA G9A were not established as fully viable stocks. It may be that these mutants are also developmentally toxic to the extent that no transgenic individuals can be produced. However, this seems unlikely due to the relative tolerance of these variants in our yeast models, as well as previous establishment of a Thr-AGT G26A mutant in *Drosophila* (Isaacson et al., 2024) which would be predicted to mistranslate more frequently than the G9A mutant attempted here.

Therefore, it may be that the lack of transgenic individuals created here stems from procedural errors in the injection protocols. This uncertainty in procedure can be extended to the tRNA mutants lacking secondary mutation discussed above, making it difficult to make definitive claims regarding the viability of these mutants. As such, no statistical comparisons are being conducted, and these data should be treated merely as guiding information for further attempts at establishing models of mutant tRNA in *Drosophila*.

Of the tRNA mutant *Drosophila* lines containing G9A secondary mutation created here, none produced any developmental phenotypes. Previous models of mistranslation have, without fail, all demonstrated some form of developmental lethality or increase in developmental time. As such, the lack of developmental phenotypes produced by tRNA mutants bearing G9A secondary mutation can be interpreted as a substantial enough reduction in mistranslation frequency such as to be tolerated effectively throughout development.

Although the tRNA mutant lines established here produce no change to developmental viability, there may be other phenotypes affected by their relatively low frequencies of mistranslation. For instance, some of the mistranslating lines, individuals that survive into adulthood display an increase in longevity, potentially through a hormetic mechanism. Such an effect could also occur for the mildly mistranslating lines created here. Hormetic effects, if present, could also extend to neural functions and affect climbing and/or learning and memory.

Given the apparent inviability of mistranslating tRNA mutants lacking secondary mutations, and the lack of phenotypes observed in those with a G9A mutation, the frequency of mistranslation produced by the combination of the G26A mutation, the tSerUGA1-1 gene, and the specific landing site used here, represents an experimental “sweet spot” or "Goldilocks’s zone”. Use of these conditions together effectively balances the frequency of mistranslation to allow viability but also permit observable biological phenotypes. A subset of tRNA variants with G26A mutations used here were also used in (Isaacson et al., 2024) to great success. It bears repeating that this “Goldilocks’s” trend for G26A variants is likely to change with the use of alternative tRNA genes or transgenic landing sites.

The hurdles of assessing mistranslating tRNA variants in *Drosophila* often stem from the problem of being unable to control the mutant tRNA’s expression; excessive expression and mistranslation can increase both inviability and likelihood of suppression. While the addition of secondary mutations mitigates these severe negative effects enough to permit experimentation, their presence, by design, dampens the observed phenotypes and may mask others entirely. As such, it may be desirable to prioritize the creation of *Drosophila* mutant mistranslating tRNA lines lacking secondary tRNA mutations. It is possible that more practiced hands may eventually produce transgenic offspring for each of the failed lines attempted here. However, given their prospective high rates of mistranslation, these lines would be especially susceptible to the emergence of suppressor mutations that ameliorate mistranslation’s effects and obfuscate their study.

For these reasons, the development of a conditional expression system for tRNAs in *Drosophila*, a la the Gal4/UAS system or the yeast doxycycline inducible system used here, would be exceptionally valuable. This approach would allow for the conditional suppression of mutant tRNA expression during development to prevent early die off. Use of a binary expression system in which the multiple components required for mutant expression are maintained as separate stocks would also aid in prevention of suppressor emergence. A binary expression system would not only solve these problems but also open the door to a huge number of new experimental opportunities, such as assessing how different forms of mistranslation affect specific cell types or particular developmental time points.

## Conclusion

The data presented here demonstrate a wide breadth of phenotypes for different forms of mistranslation. Several characteristics of mutant mistranslating tRNAs are identified as contributing to the phenotypes produced, such as the severity of biochemical difference in substituted amino acids, frequency of mistranslation, the synonymous codons mistranslated, and the identity of the specific tRNA gene bearing the mistranslating mutation. These experiments also demonstrate the utility of using *Saccharomyces cerevisiae* as a model for *Drosophila* tRNA mutant mistranslation, such that mutants can be assessed prior to laborious attempts at *Drosophila* model reaction.

## Data Availability Statement

Fly lines and plasmids are available upon request. The authors affirm that all data necessary for confirming the conclusions of the article are present within the article, figures, and Supplemental material. Supplementary Figures S1-S2, Supplementary Methods, and Supplementary Tables S1- S3 are appended to the manuscript.

## Acknowledgements

We would like to thank Dr. Chris Brandl, Dr. Matthew Berg, and Julie Genereaux for their invaluable insight and donation of plasmids and yeast strains. We would also like to thank Dr. Kyle Hoffman and Bioinformatics Solutions for their efforts in collecting and analyzing the mass spectrometry experiments.

## Generative AI Statement

No generative AI was used in this manuscript.

## Supplemental Methods: Mass Spectrometry

***In solution digestion*:** Yeast cell pellets were resuspended Y-PER (ThermoFisher Scientific) with Halt™ Protease and Phosphatase Inhibitor Cocktail (ThermoFisher Scientific). Each sample was sonicated at an amplitude of 60% for 5min with cycles of 30s on/30 off. Cell debris was pellet, and soluble protein was quantified by using Pierce™ BCA Protein Assay kit. An aliquot of 10µg per sample was the reduced with 100mM tris(2-carboxyethyl)phosphine (TCEP) and alkylated with 200mM IAA. The lysate was then digested with trypsin (New England Biolabs) at a 50:1 (protein:enzyme) ratio according to manufacturer’s recommendations. Digested peptides were desalted by C18 clean-up, lyophilized, and stored until analysis.

***Mass spectrometry (MS/MS) analysis*:** The digested peptides were resuspended in buffer A (0.1% FA) and injected into the timsTOF Pro (Bruker Daltronics, Bremen, Germany) by nanoflow liquid chromatography using a Thermo Scientific EASY-nLC 1000 (ThermoFisher, Massachusetts, USA). Liquid chromatography was performed using a constant flow of 300 µl/min and a 15cm reversed-phased column with a 75 µm inner diameter filled with Reprosil C18 (PepSEP, Bruker, Germany). Mobile phase A was 0.1% Formic Acid and Mobile phase B was 99.9% Acetonitrile, 0.1% Formic Acid. Peptide separation was carried out over 55.5 minutes as follows; linearly 4% A to 35% B over 50 minutes with an increase to 100% B over 30 seconds and held constant for 5 minutes to clean the column. Column equilibration was done prior to sample loading automatically.

The *tims*TOF Pro was outfitted with a Captivespray source (Bruker Daltronics, Bremen, Germany), operated in PASEF mode. Trapped ion mobility separation was achieved by using an accumulation time of 100ms in the first TIMS region and ramps of the TIMS region from 0.85 V^.^s/cm^2^ to 1.30 V^.^s/cm^2^, with each ramp lasting 100ms. MS and MS/MS scans were limited to 100 m/z to 1700 m/z, and a polygon filter was applied to the m/z and ion mobility dimensions to select for multiple charged ions most likely to be peptide precursors. Collision energy was applied as a function of ion mobility with a linear regression using the following parameter settings: 0.85 V^.^s/cm^2^ ◊ 27eV, 1.30 V^.^s/cm^2^ ◊ 45eV. TIMS voltage was calibrated using ions from the Agilent Tune Mix (m/z 622, 922, 1222). Active Exclusion of MS/MS scans were enabled at a setting of 0.40min. Quadrupole isolation was set to 2m/z for m/z less than 700, and 3.0 m/z for ions with an m/z greater than 800. A linear regression calculation was done automatically for ions in between m/z 700 and 800.

MS Raw Files were processed using PEAKS Studio 13 (Bioinformatics Solutions Inc., Ontario, Canada). The data was searched against the reviewed *Saccharomyces cerevisiae* database (Uniprot) with a PSM score threshold of -10lgP ≥ 20. Precursor ion mass error tolerance was set to 20 ppm and fragment ion mass error tolerance was set to 0.05 Da. Semi-specific cleavage with trypsin was selected with a maximum of 2 missed cleavages. A fixed modification of carbamidomethylation (+57.02 Da) on cysteines and variable modifications of deamidation (+0.98 Da) on asparagine and glutamine, as well as oxidation (+15.99 Da) on methionine, were specified. Mis-incorporation of Ser was identified by searching for fragment ion mass shifts of +30.01 Da at Gly, -26.05 Da at Ile, and -12.04 Da at Val positions. All reported mutations pass confident modification parameters set in PEAKS Studio, including peptide tag length ≥ 3 and peptide tag relative ion intensity ≥ 2%. A 1% false discovery rate was set for the database search.

---

**Supplementary Figure S1.**
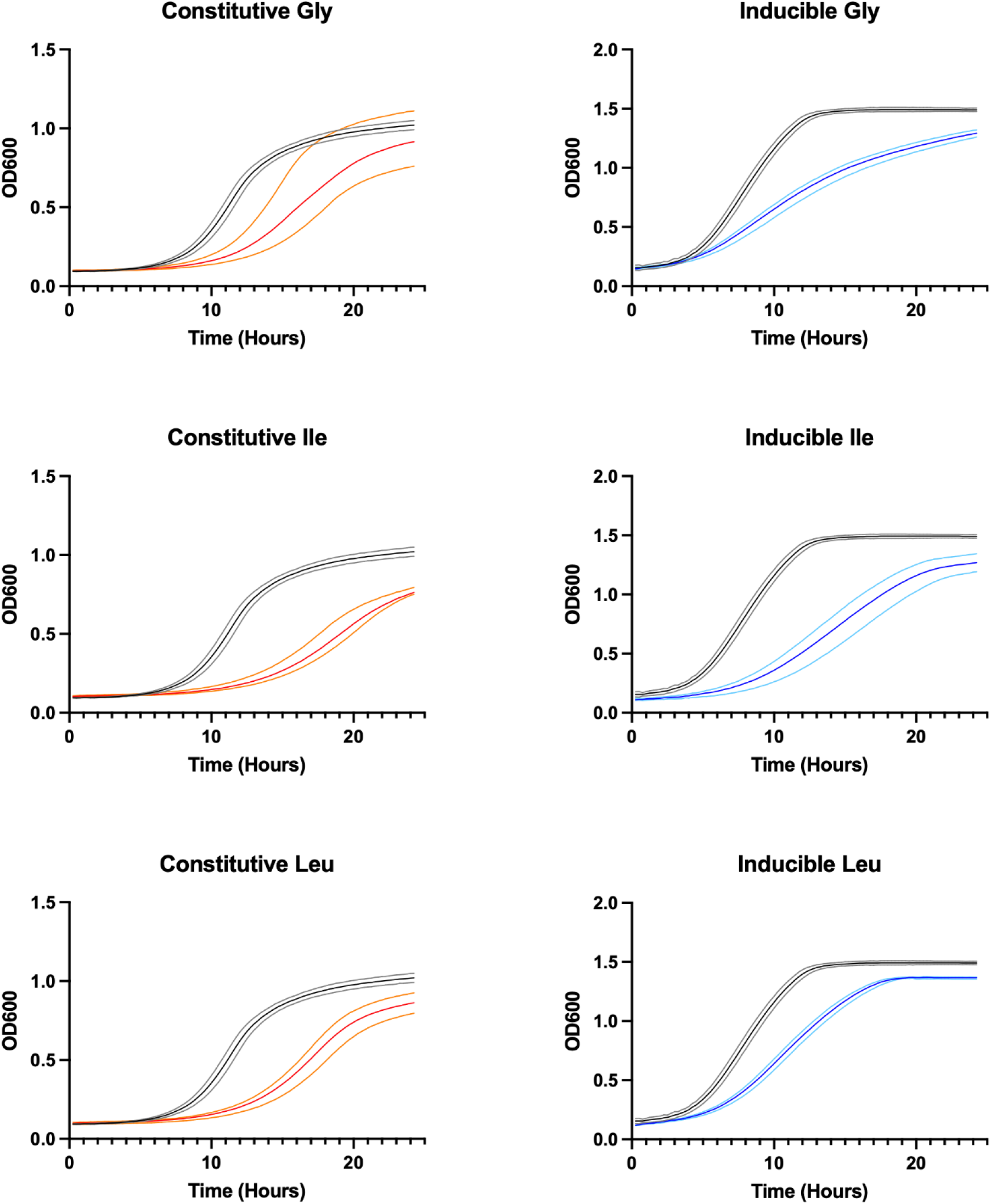
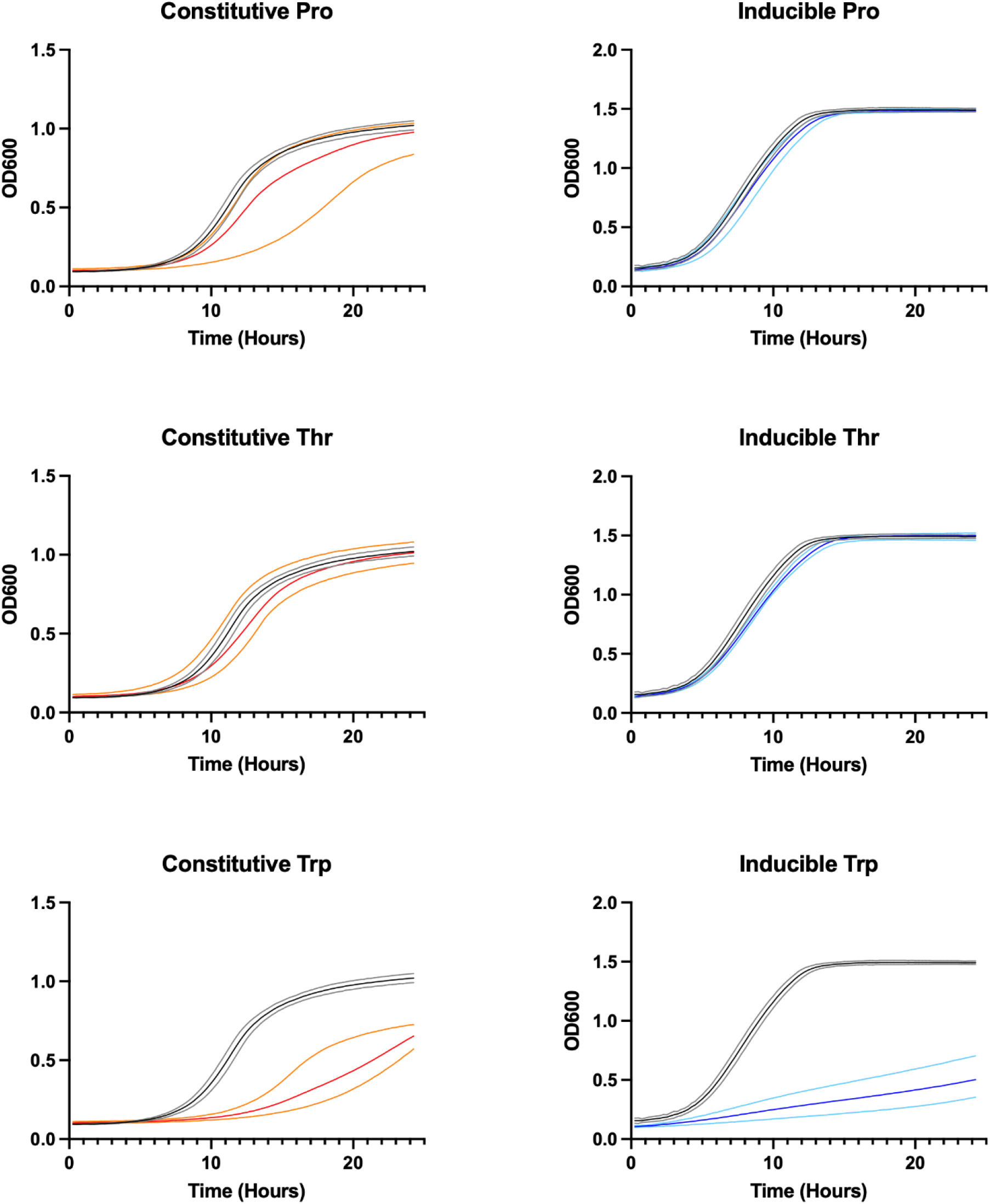
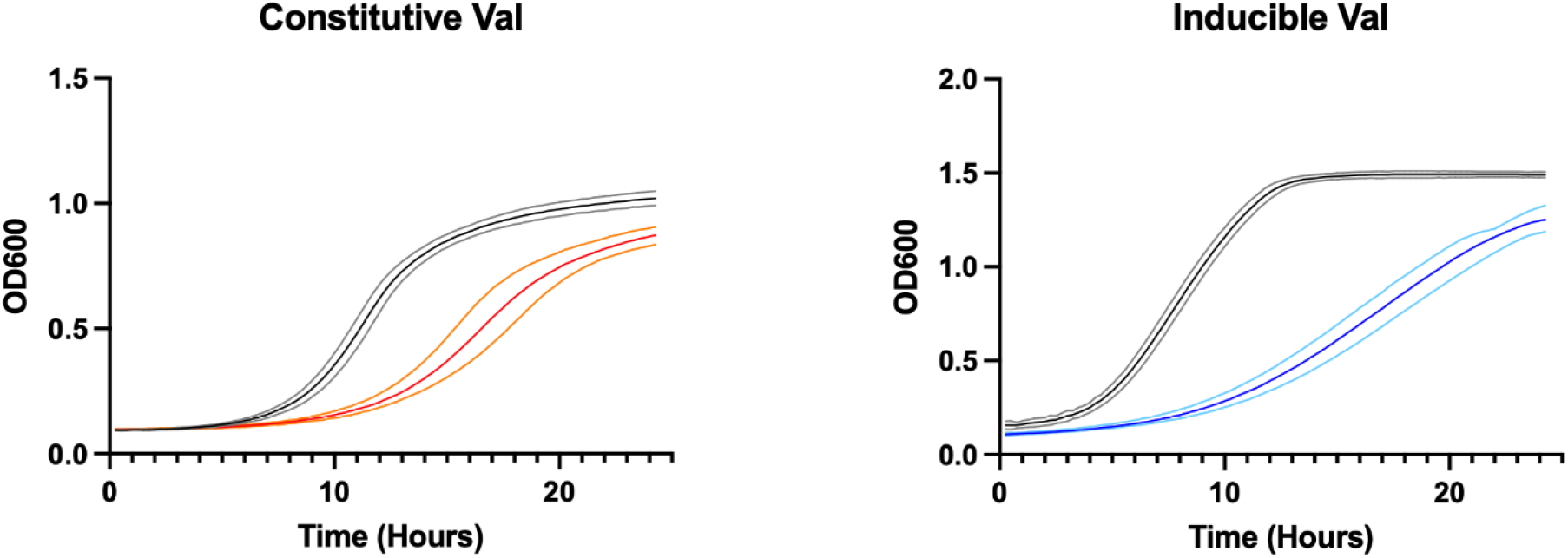
– Growth curves of mistranslating tRNA variants summarized in. Figure 1. Black lines denote average growth of control yeast expressing *D. mel* tSerUGA1-1, while gray lines denote standard deviation among triplicates. Left: red lines denote growth of cells constitutively expressing mutant tRNAs, while orange lines denote standard deviation among triplicates. Right: blue lines denote growth of yeast cells inducibly expressing mutant tRNAs via doxycycline inducible tRNA expression system, while lighter blue lines denote standard deviation among triplicates.

**Supplementary Figure S2.**
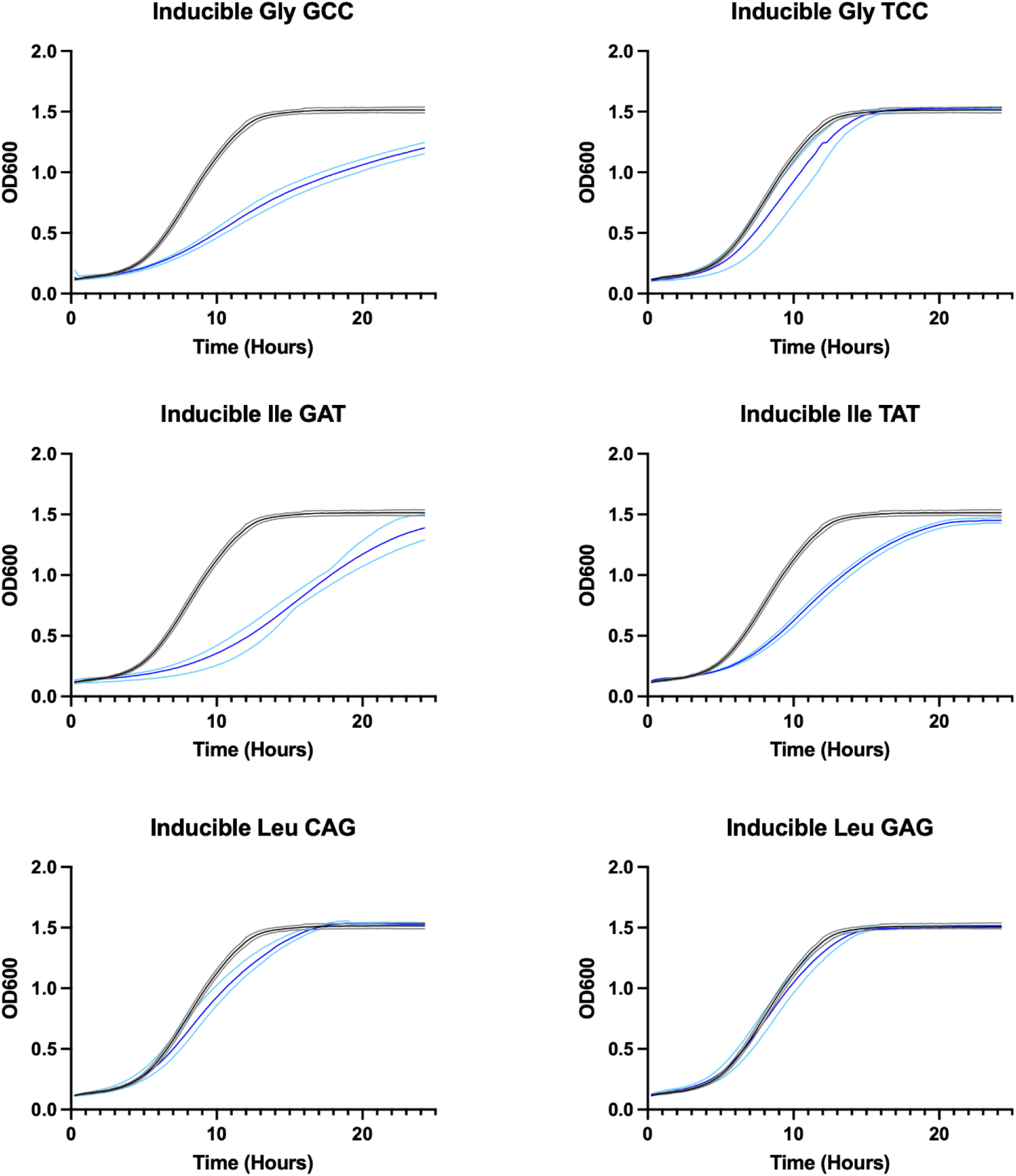
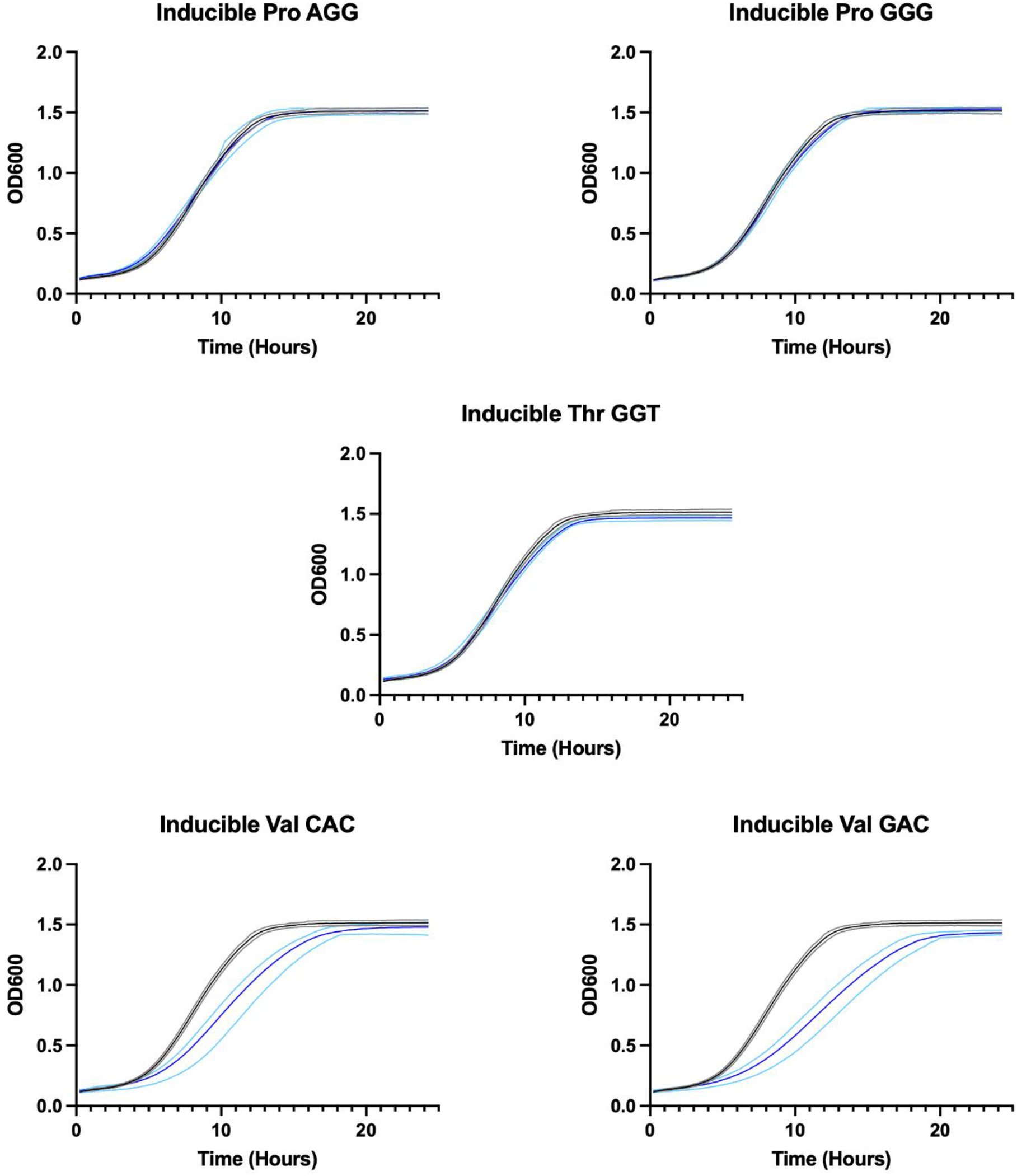
– Growth curves of mistranslating tRNA variants summarized in. Figure 3. Black lines denote average growth of control yeast expressing *D. mel* tSerUGA1-1, while gray lines denote standard deviation among triplicates. Blue lines denote growth of yeast cells inducibly expressing mutant tRNAs via doxycycline inducible tRNA expression system, while lighter blue lines denote standard deviation among triplicates.

